# Brain-wide distributed processing underlying natural vision and audition

**DOI:** 10.64898/2026.05.26.727967

**Authors:** Marco D’Alessandro, Antonino Greco, Francesca Setti, Emiliano Ricciardi, Giovanni Pezzulo

## Abstract

A key challenge in neuroscience is understanding how large-scale brain dynamics give rise to perception under naturalistic conditions. Here we address this question using multivariate information decomposition of functional interactions between brain regions while typically developed, congenitally blind and congenitally deaf participants experienced audiovisual, auditory-only or visual-only versions of the same narrative. This approach allowed us to resolve information dynamics into redundant components, indexing shared information and sensory robustness, and synergistic components, indexing distributed processing that emerges only through joint interactions between brain regions. Redundant interactions were closely aligned with conventional functional connectivity and captured sensory modality- and experience-dependent differences mainly within sensory systems, whereas synergistic interactions captured these differences across high-level cortical areas. Crucially, prefrontal cortex coordinated brain dynamics through synergistic interactions independently of sensory modality and previous sensory experience, consistent with a modality-independent hub that encodes high-level semantic information and orchestrates the demands of naturalistic perception. Moreover, multimodal perception elicited the strongest high-order synergistic interactions across distributed brain subsystems, followed by unimodal visual and auditory perception, whereas sensory deprivation reduced this distributed architecture. Finally, synergistic and redundant interactions jointly supported the encoding of low-level sensory and high-level perceptual features, revealing how the trade-off between sensory robustness and distributed processing is organized across the human brain. Together, these results reveal a brain-wide information architecture for natural vision and audition, illuminating how distributed processing in the human brain supports rich perceptual experience.

## Introduction

Perception in real-world settings requires the brain to transform continuous, multimodal sensory input into coherent interpretations of the environment. Understanding how this transformation emerges from large-scale brain dynamics remains a central challenge in neuroscience ^1–4^.

Classical accounts often describe perception as a sequence of computations progressing from modality-specific sensory cortices toward high-level transmodal areas ^5–8^. However, this view is increasingly complemented by evidence that cognition and perception rely on brain-wide dynamics, with representations and computations distributed across multiple interacting regions ^9–13^. Recent large-scale recordings in animals and humans have emphasized that complex behaviour engages widespread neural populations rather than isolated processing modules ^14–25^, raising key questions about how distributed information processing supports perception and whether specific cortical hubs coordinate these brain-wide interactions.

Naturalistic paradigms provide a powerful way to address these questions ^26–30^. Movies and narratives are particularly well-suited because they preserve the full hierarchy of perceptual structure present in real-world experience: from low-level, modality-specific sensory dynamics, such as the spatio-temporal statistics of visual motion and the spectral properties of acoustic input, to high-level features including object and scene identity, and up to semantic and narrative content that unfolds over extended timescales ^31–34,30^. They therefore make it possible to study distributed brain dynamics during the continuous integration and interpretation of sensory input, context and meaning. Moreover, the same narrative can be presented through different sensory formats and to individuals with different sensory histories, providing a unique opportunity to dissociate modality-specific sensory processing, lifelong sensory experience, and modality-(in)dependent representations of perceptual content.

To characterize distributed processing in the brain, previous studies have often relied on multivariate decoding ^35^ and functional connectivity ^36^. Decoding approaches reveal where information about stimuli or behaviour can be recovered across the cortex ^37–40^, whereas functional connectivity measures statistical coupling among brain regions ^41–46^. However, neither approach fully specifies the informational structure of distributed processing. Widespread decodability does not determine whether information is duplicated across regions or depends on their joint activity. Similarly, functional connectivity reveals coupling, but not whether this coupling reflects shared or integrated information, whether it encodes specific perceptual features of the stimulus, or whether it extends beyond pairwise interactions.

Multivariate information decomposition methods offer a way to make these distinctions explicit ^47–56^. Partial Information Decomposition (PID) ^57–59^ separates the information that multiple sources provide about a target into distinct informational atoms, allowing *redundant* information shared across sources to be distinguished from *synergistic* information available only through their joint state. Integrated Information Decomposition (IID) extends this logic to dynamical systems, decomposing how the past state of a system constrains its future state into distinct redundant and synergistic components ^47,53,54^. In parallel, high-order interaction (HOI) measures such as O-information quantify whether multivariate systems are dominated by redundancy or synergy across interactions involving more than two brain regions ^60–64^. Together, these methods provide an information-resolved framework for studying distributed processing, linking brain-wide interactions to the form, scale and content of the information they carry.

Crucially, redundancy and synergy capture complementary aspects of distributed processing. Redundancy reflects information that is available across multiple regions and is therefore distributed in the sense that the same information is present in different parts of the brain. This distributed availability may support sensory robustness, because information does not depend on a single cortical site. Synergy provides a stricter definition of distributed processing, as it captures information that is not available in any region alone but emerges only when multiple regions are considered jointly. In this sense, synergy identifies cases in which information is genuinely interaction-dependent, requiring the integration of partial signals across distributed brain areas. Thus, whereas redundancy reveals how information is spread across the brain, synergy reveals when information depends on integration across regions.

Here, we used an information-theoretic framework, based upon the notions of redundancy and synergy, to characterize brain-wide information dynamics during natural vision and audition. We analyzed fMRI responses while typically developed, congenitally blind and congenitally deaf participants experienced audiovisual, auditory-only or visual-only versions of the same movie narrative: the *101 Dalmatians* ^65,66^. This design assesses which aspects of brain-wide information dynamics depend on current sensory richness, which depend on lifelong sensory experience, and which are preserved across all experimental groups. To address these questions, we first used conventional functional connectivity to establish a baseline description of large-scale coupling. We then applied three complementary information-decomposition analyses. IID ^53^ separated pairwise temporal interactions into persistent redundancy and persistent synergy. HOI analysis ^64^ tested whether synergistic structure extended beyond pairs into multiregional brain subsystems. Finally, PID ^57^ quantified how pairs of regions redundantly or synergistically encoded low-level sensory and high-level perceptual features of the movie.

## Results

### Experimental design and functional connectivity

We analyzed whole-brain fMRI responses during presentation of the same naturalistic movie narrative in participants with typical development (TD) and in participants with congenital sensory deprivation (SD). Typically developed individuals viewed the movie in audiovisual, visual-only, or auditory-only format, whereas congenitally deaf and congenitally blind participants experienced the corresponding unimodal versions (Fig. 1A).

**Figure 1.**
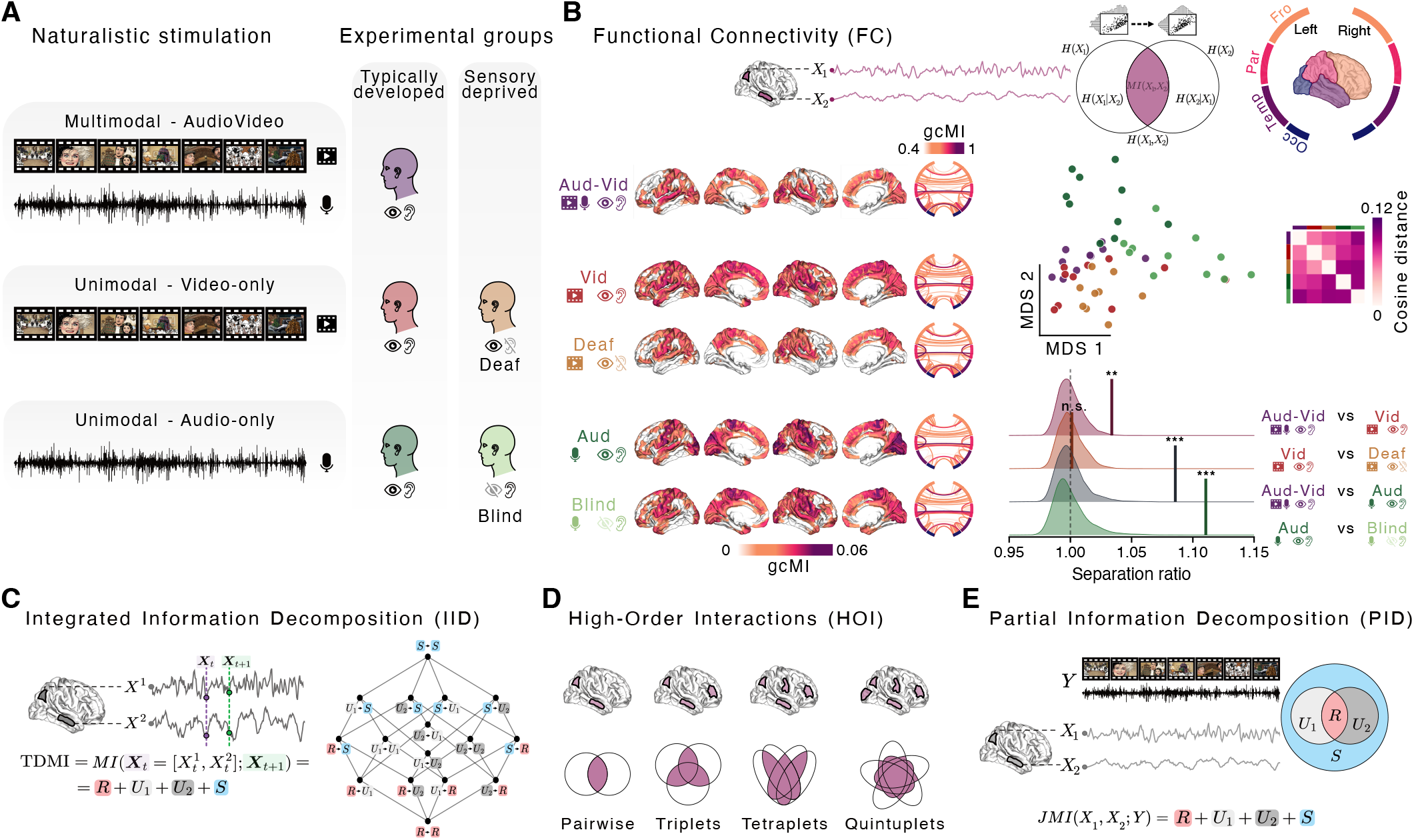
Experimental design and information decomposition analysis outline of brain-wide dynamics. **A.** Participants experienced the same naturalistic movie narrative inside the fMRI scanner under audiovisual, visual-only or auditory-only stimulation. Typically developed (TD) participants completed all three multimodal and unimodal stimulation conditions, whereas sensory deprived (SD) participants were probed with the matched unimodal conditions: visual-only for congenitally deaf and auditory-only for congenitally blind participants. This design allowed us to study the effect of multimodal versus unimodal stimulation, and the effect of sensory experience by comparing TD and SD participants under matched unimodal input. **B**. Pairwise functional connectivity was computed across cortical regions using the Gaussian-copula mutual information (gcMI) estimator. Left, circular graph plots showing group-average functional coupling patterns for the top 100 connections in each experimental group, while brain maps show the betweenness centrality of each cortical region. Right, multidimensional scaling (MDS) provides a two-dimensional projection of the cosine-distance matrix computed from participant-level connectivity profiles. Each dot represents one participant and is color-coded by experimental group. The inset shows the cosine-distance matrix averaged by experimental group. Separation-ratio plots show the observed statistic for each experimental contrast against the corresponding null distribution obtained by permutation testing. Asterisks indicate statistical significance, * P<0.05, ** P<0.01, *** P<0.001; n.s., not significant. **C-E**. Schematic of the information decomposition framework used in subsequent analyses. Integrated Information Decomposition (IID) decomposed the temporal delayed mutual information (TDMI) to quantify how much information about future brain dynamics trajectory is carried redundantly or synergistically by pairs of brain regions. High-order interactions (HOI) analysis identified maximally synergistic interactions across multiregional brain subsystems. Partial Information Decomposition (PID) quantified redundant and synergistic encoding of perceptual stimulus features. Together, these information-resolved analyses move beyond conventional functional connectivity to characterize how sensory modality and sensory experience shape distributed information dynamics during natural vision and audition. Due to copyright, the original movie frames have been replaced with images generated using deep image generative models.

This experimental design allowed us to examine how brain-wide naturalistic perception is shaped by the richness of current sensory input and the sensory history through which the brain has developed. Specifically, it allowed us to identify aspects of brain-wide information dynamics that vary with current sensory richness or lifelong sensory experience, and to reveal patterns shared across all experimental groups, independent of both sensory modality and previous sensory experience.

As a first step, we characterized brain-wide interactions underlying naturalistic perception using a conventional functional connectivity approach (Fig. 1B left). This established a reference point for characterizing large-scale brain dynamics using a classical measure of interregional coupling ^36^. For each participant, we estimated pairwise functional connectivity across cortical regions using Gaussian-copula mutual information (gcMI) ^67^, compared whole-brain connectivity profiles using cosine distance, and projected the resulting distance matrix into a two-dimensional space using multidimensional scaling (MDS). Connectivity patterns revealed widespread large-scale interactions across brain areas in all experimental groups, with distributed cortical hubs spanning sensory and associative transmodal regions. The resulting geometry showed a structured organization of the experimental groups, with participant connectivity profiles clustering according to stimulation condition and sensory experience.

To quantify this organization, we computed a separation ratio (SR) for each contrast, defined as the ratio between between-group distance and within-group distance (Fig. 1B right). We tested four principal contrasts of interest. Comparisons between audiovisual (Aud-Vid) and video-only (Vid), and between audiovisual and audio-only (Aud), isolate the contribution of sensory richness within TD participants (multimodality vs. unimodality). Comparisons between video-only TD participants and congenitally deaf participants (Deaf), and between audio-only TD participants and congenitally blind participants (Blind), instead probe the impact of sensory developmental condition under matched unimodal input (TD vs. SD). Results showed that the multimodal condition was reliably separated from both audio-only (SR = 1.09, P < 0.001) and video-only (SR = 1.03, P = 0.002) conditions. Under matched unimodal input, TD participants were clearly separated from blind participants in the audio-only condition (SR = 1.11, P < 0.001), but not from deaf participants in the video-only condition (SR = 1.00, P = 0.406).

Although conventional functional connectivity provides a compact description of large-scale coupling, it cannot determine the informational structure of these functional interactions. It can show that distant brain regions have similar temporal course activity, but not whether this reflects redundant information available in multiple regions or synergistic information indexing distributed processing ^47^. Nor can it determine whether distributed neural interactions extend beyond pairs of regions, or what stimulus features these interactions encode. These distinctions are central to understanding how distributed information processing underlying perception is organized.

We therefore used multivariate information decomposition to move from measuring coupling between brain regions to characterizing the informational structure and the distributed nature of their interactions. To achieve this, we took three complementary approaches. Integrated Information Decomposition (IID) ^53^ separated pairwise temporal interactions into persistent redundancy and persistent synergy (Fig. 1C). High-order interaction analysis ^65^ tested whether synergistic structure extended beyond pairs into multiregional brain subsystems (Fig. 1D). Finally, Partial Information Decomposition (PID) ^57–59^ quantified how pairs of regions redundantly or synergistically encoded low-level sensory and high-level perceptual features of the movie (Fig. 1E).

### Brain-wide temporally persistent redundant and synergistic interactions

We applied IID and quantified, for each pair of regions, persistent redundancy and persistent synergy (Fig. 2A). Persistent redundancy captures information about future joint activity that is duplicated across two regions, whereas persistent synergy captures information about the future that is available only from the two regions considered together ^53^.

**Figure 2.**
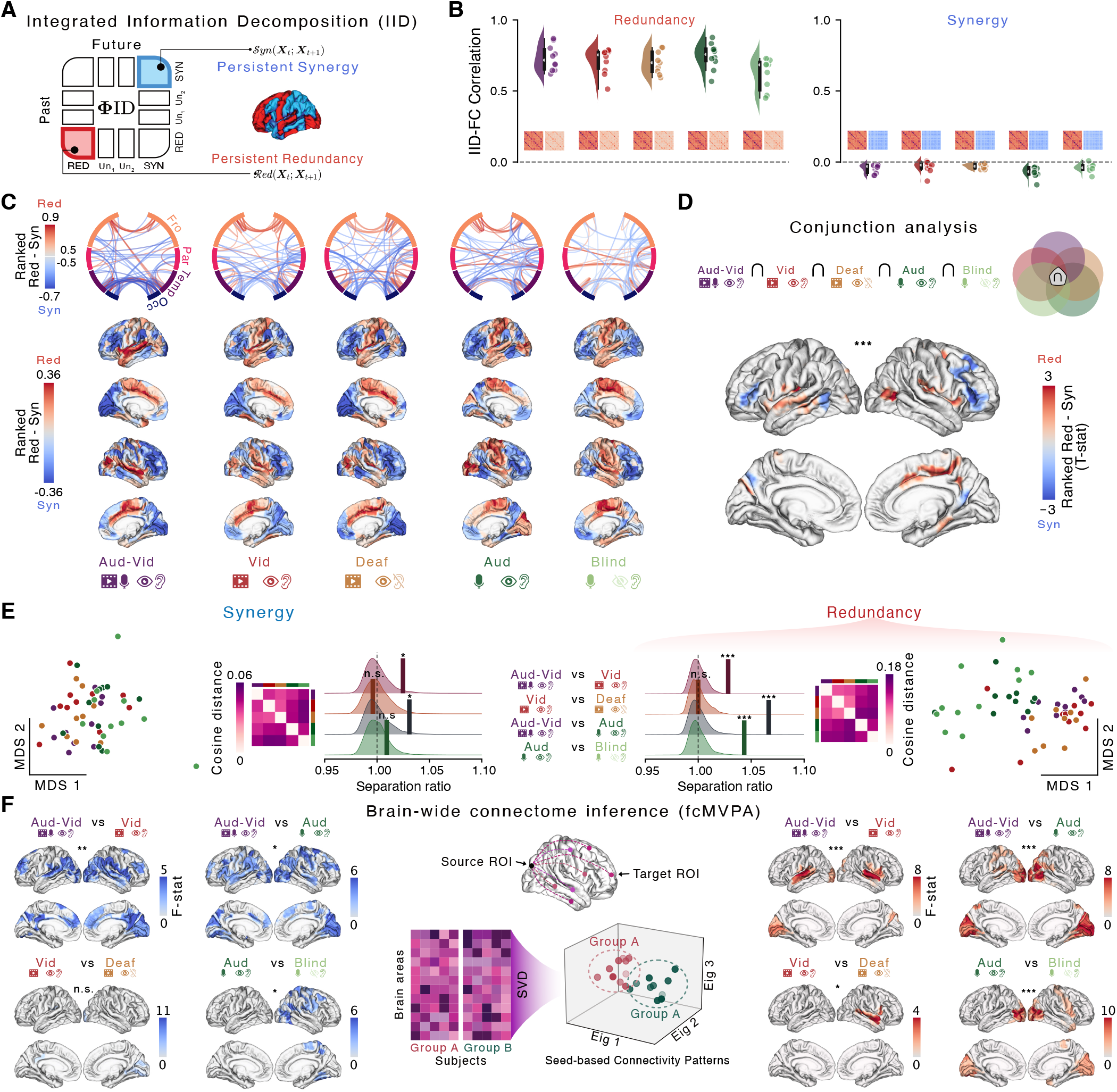
Integrated Information Decomposition reveals condition-specific and invariant brain-wide redundant and synergistic interactions. **A.** Schematic illustration of the Integrated Information Decomposition (IID) analysis. For each pair of cortical regions, the information that their past state provides about their future activity over the course of the movie was decomposed into distinct informational atoms. We focused on temporally persistent redundancy, indexing information maintained in common across brain areas, and temporally persistent synergy, indexing information maintained only through their joint dynamics. **B**. Pearson correlation coefficients between functional connectivity profiles and redundant or synergistic interaction profiles. Each dot represents individual participants, while insets show group-average connectivity matrices and interaction matrices for redundancy and synergy. **C**. Circular plots showing group-average ranked redundancy minus synergy scores for the top 100 interactions (50 per information atom), while brain maps show regional betweenness centrality along this redundancy-to-synergy gradient. **D**. Conjunction analysis identifying redundancy-synergy gradients common to all experimental groups, revealing interaction cortical hubs invariant to sensory modality and previous sensory experience. Brain maps show conjunction t-statistics, with asterisks and transparency indicates statistical significance determined by cluster correction and permutation testing. **E**. Multidimensional scaling (MDS) projections of cosine-distance matrices computed from redundancy and synergy interaction profiles. Each dot represents one participant, color-coded by group, while insets show experimental condition-averaged distance matrices. Separation-ratio plots show observed contrast statistics against permutation null distributions. **F**. Brain-wide connectome inference of synergistic and redundant interactions. For each seed region, whole-brain interaction profiles were compared across experimental contrasts using functional connectivity multivariate pattern analysis (fcMVPA). Brain maps show cortical regions whose interaction patterns differed between conditions, with asterisks and transparency indicates statistical significance. Legend: * P<0.05, ** P<0.01, *** P<0.001; n.s., not significant.

We first assessed whether the IID results aligned with those obtained using conventional functional connectivity, and whether the latter primarily reflected redundant or synergistic interactions. For this, participant-level functional connectivity profiles were correlated with the corresponding redundancy and synergy interaction profiles derived from IID (Fig. 2B). Across all experimental groups, redundancy showed strong correspondence with conventional connectivity (all P < 0.0001, mean and range *r* = 0.70 [0.62, 0.75]), whereas synergy did not (all P > 0.05, mean and range *r* = -0.04 [-0.03, -0.07]). These results indicate that standard functional connectivity mainly captures temporally persistent information shared across regions, while synergistic interactions capture distinct forms of neural interactions beyond coupling strength alone.

We next examined the large-scale cortical organization of temporally persistent redundancy and synergy (Fig. S1) by ranking interactions according to their redundancy-to-synergy balance and identifying their cortical hubs (Fig. 2C). Across all experimental groups, redundancy-dominated interactions were expressed more strongly within modality-specific temporal and parietal regions, whereas synergy-dominated interactions were more prominent across fronto-parietal transmodal cortices. In conditions involving visual stimulation, synergistic hubs further extended into occipital regions. Statistical maps confirming the significant cortical distribution of these redundancy-to-synergy gradients within each experimental group are shown in Fig. S2A (all P < 0.037 cluster-corrected).

At this point, we could directly address one of the central questions of this study: are some aspects of brain-wide information dynamics preserved independently of current sensory modality and previous sensory experience? To test this, we performed a conjunction analysis, identifying cortical regions that consistently occupied similar positions along the redundancy-to-synergy gradient across all experimental groups, using a robust statistical permutation framework. Crucially, we indeed identified invariant redundant and synergistic cortical hubs (Fig. 2D).

Redundancy-dominated regions were consistently concentrated within temporal and parietal cortices (all clusters P < 0.045 cluster-corrected), whereas synergy-dominated regions were distributed across frontal and parieto-occipital areas (all clusters P < 0.0002 cluster-corrected). To further confirm the validity of these findings, we repeated the conjunction analysis by intersecting the group-wise significant redundant and synergistic cortical regions, revealing a highly similar cortical distribution, particularly for the synergistic cortical hubs in prefrontal cortex (Fig. S2B). Together, these results suggest that large-scale information dynamics during naturalistic perception are organized around invariant cortical hubs, with the prefrontal cortex emerging as a modality-independent hub orchestrating distributed processing through brain-wide synergistic interactions.

Having identified invariant cortical architectures shared across all experimental groups, we next asked which aspects of redundant and synergistic interactions varied as a function of current sensory richness or previous sensory experience. To address this, we firstly compared whole-brain interaction profiles using the same separation ratio and MDS approach previously applied to conventional functional connectivity (Fig. 2E). Redundant interactions largely recapitulated the organization previously observed for functional connectivity, showing reliable separation both as a function of sensory richness (Aud-Vid vs Aud: SR = 1.07, P < 0.002; Aud-Vid vs Vid: SR = 1.03, P = 0.002) and, for the matched auditory comparison, sensory experience (Aud vs Blind: SR = 1.03, P < 0.001; Vid vs Deaf: SR = 1.00, P = 0.437).

By contrast, synergistic interaction profiles showed a different organizational pattern. Although reliable differences still emerged as a function of sensory richness (Aud-Vid vs Aud: SR = 1.03, P = 0.020; Aud-Vid vs Vid: SR = 1.02, P = 0.033), these effects were consistently smaller than those observed for redundancy, while differences associated with lifelong sensory experience were no longer significant (Aud vs Blind: SR = 1.01, P = 0.165; Vid vs Deaf: SR = 0.99, P = 0.611). Together, these results suggest that whole-brain synergistic interactions were comparatively more similar across different sensory modalities and lifelong sensory histories, consistent with the idea that these interactions may reflect more modality-independent aspects of perceptual processing. To localize the cortical networks driving condition-dependent effects, we next performed a brain-wide connectome inference analysis using functional connectivity multivariate pattern analysis (fcMVPA). This approach tests whether the brain-wide interaction patterns of a given cortical region differ across experimental conditions, allowing changes in multivariate patterns of redundant and synergistic interactions to be mapped at the regional level (Fig. 2F). Results revealed strikingly different spatial organizations for redundancy and synergy. Redundant interactions largely followed modality-specific sensory differences, with both multimodal versus unimodal contrasts and sensory deprivation effects expressed predominantly within cortical areas corresponding to the absent sensory modality. In particular, conditions lacking auditory input primarily involved temporal cortices (Aud-Vid vs Vid: all clusters P < 0.019 cluster-corrected, all ηp^2^ > 0.51; Vid vs Deaf: P = 0.044 cluster-corrected, ηp^2^ = 0.57), whereas conditions lacking visual input mainly involved occipital and parietal regions (Aud-Vid vs Aud: P < 0.001 cluster-corrected, ηp^2^ = 0.59; Aud vs Blind: all clusters P < 0.016 cluster-corrected, all ηp^2^ > 0.50). Of note, deprived unimodal areas in both Blind and Deaf showed different patterns of redundant interactions.

Crucially, synergistic interaction patterns instead showed broader condition-dependent differences, involving large-scale networks spanning both sensory and high-level associative regions. In particular, multimodal contrasts revealed widespread effects across all cerebral lobes (Aud-Vid vs Vid: P = 0.008 cluster-corrected, ηp^2^ = 0.57; Aud-Vid vs Aud: P = 0.013 cluster-corrected, ηp^2^ = 0.55), while sensory deprivation effects were only observed when comparing blind participants with TD participants under auditory-only stimulation, primarily within parieto-occipital regions (Aud vs Blind: all clusters P < 0.037 cluster-corrected, all ηp^2^ > 0.52; Vid vs Deaf: P > 0.05 cluster-corrected).

Taken together, these IID results reveal that perceptual processing relies on two complementary forms of distributed brain organization. Redundant interactions primarily reflect the sensory structure of the stimulus itself, changing according to which sensory information is available. Synergistic interactions instead reveal a more stable and distributed architecture that is comparatively preserved across sensory modalities and previous sensory experience. Most notably, prefrontal cortex consistently emerged as a central hub of these distributed synergistic interactions across all experimental conditions, suggesting a role in orchestrating the cognitive and attentional demands of naturalistic perception, regardless of the modality involved. Importantly, redundant interactions largely aligned with conventional functional connectivity approaches, but synergistic interactions diverged significantly from them, highlighting how information decomposition can uncover the informational architecture underlying large-scale brain dynamics during naturalistic perception.

### Brain-wide synergistic high-order interactions

The widespread and comparatively modality-independent synergistic interactions identified by the previous analyses revealed distributed cortical processes spanning large-scale brain networks. We therefore sought to characterize these dynamics at the level of high-order brain interactions, namely statistical interactions emerging from the coordinated activity of more than two brain regions simultaneously ^68^. Such HOI are increasingly recognized as a fundamental feature of complex systems, reflecting collective dynamics that cannot be reduced to simpler pairwise relationships ^69^. We therefore used high-order interaction analysis to identify and characterize maximally synergistic multiregional cortical subsystems engaged during naturalistic perception (Fig. 3A).

**Figure 3.**
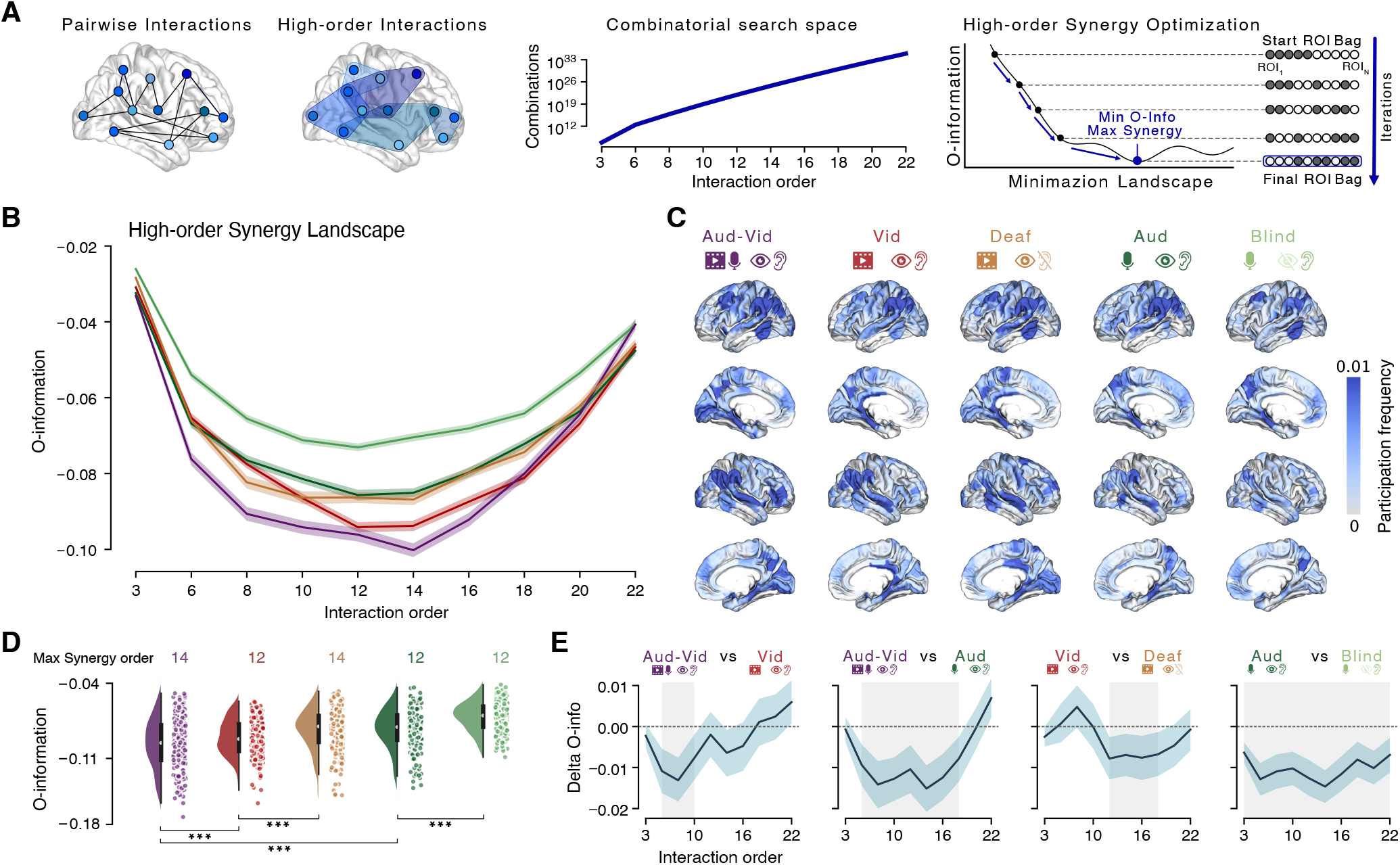
Identifying maximally synergistic brain subsystems with high-order interactions. **A.** Schematic illustration of the high-order interaction (HOI) analysis, depicting pairwise interactions contrasted with high-order interactions emerging across multiregional brain subsystems. The combinatorial growth of the search space across the cortical parcellation illustrates the large number of possible region combinations at increasing interaction order. Simulated annealing was used to efficiently explore this space and identify brain subsystems with maximal high-order synergy using O-information to quantify the balance between redundancy and synergy across high-order interactions, with more negative values indicating stronger synergy. **B**. High-order synergy landscape showing O-information as a function of interaction order for each experimental group. Shaded areas indicate standard error of the mean (SEM) across seed initializations of the annealing optimization procedure. **C**. Brain maps of participation frequency in high-order synergistic subsystems across all interaction orders, weighted by the corresponding O-information values. **D**. Experimental group comparisons at the optimal interaction order. Each dot represents seed initializations, while asterisks statistical significance (*** P<0.001). **E**. Contrast plots of the difference in O-information across interaction orders for the relevant experimental comparisons, with shaded areas indicating SEM across seed initializations. Vertical grey bars indicate statistical significance (P<0.05, cluster-corrected).

To characterize these high-order interactions, we used O-information ^60^, a multivariate information-theoretic measure that quantifies whether a system is dominated by redundant or synergistic interactions. Positive O-information indicates predominantly redundant dependencies, while negative O-information indicates predominantly synergistic interactions emerging only through the collective activity of multiple brain regions simultaneously. Because the number of possible multiregional combinations grows combinatorially with interaction order, exhaustive exploration of all subsets rapidly becomes computationally intractable (Fig. 3A). We therefore combined O-information with a simulated annealing optimization procedure to efficiently identify cortical subsystems expressing maximal high-order synergy, corresponding to subsets with maximally negative O-information value ^64^.

Across interaction orders ranging from 3 to 22 regions, all experimental groups showed progressively stronger synergistic organization with increasing subsystem size, reaching maximal synergy at intermediate interaction orders before decreasing thereafter, thus defining a characteristic high-order synergy landscape for perceptual processing (Fig. 3B). The cortical contribution to these maximally synergistic subsystems was not spatially uniform. Participation frequency brain maps revealed that high-order synergistic interactions predominantly involved distributed associative regions spanning frontal, temporal and parietal cortices, including supramarginal regions (Fig. 3C).

Across all experimental groups, maximal synergy consistently emerged at interaction orders between 12 and 14 regions (Fig. 3D). Comparing the experimental groups on these maximally synergistic subsystems across independent optimization runs revealed significant effects of both sensory richness and sensory experience. Audiovisual stimulation produced stronger high-order synergy than both visual-only (P = 0.007, Cohen’s d = 0.27) and auditory-only conditions (P < 0.001, d = 0.63), whereas TD participants showed stronger high-order synergy than both blind (P < 0.001, d = 0.72) and deaf (P < 0.001, d = 0.37) participants under matched unimodal stimulation. These effects were similarly observed when comparing the full high-order synergy landscape across all interaction orders (Fig. 3E), revealing significant effects of both sensory richness (both P < 0.001 cluster-corrected, both mean cluster d > 0.49) and sensory experience (both P < 0.001 cluster-corrected, both mean cluster d > 0.36).

Taken together, these results reveal that naturalistic perception engages large-scale cortical subsystems interacting synergistically through the integration of information across multiple brain regions. These high-order interactions were strongest during multimodal perception and were reduced by sensory deprivation under matched sensory stimulation, revealing a combined effect of both sensory richness and lifelong sensory experience.

### Brain-wide distributed processing of stimulus perceptual features

Finally, we sought to characterize the informational content carried by distributed neural interactions during naturalistic perception, providing a complementary perspective to the previous analyses of large-scale information dynamics. To address this, we used PID to decompose the joint mutual information (JMI) between pairs of cortical regions and external perceptual features, encoding both low-level and high-level stimulus content, into redundant and synergistic informational components (Fig. 4A, center).

**Figure 4.**
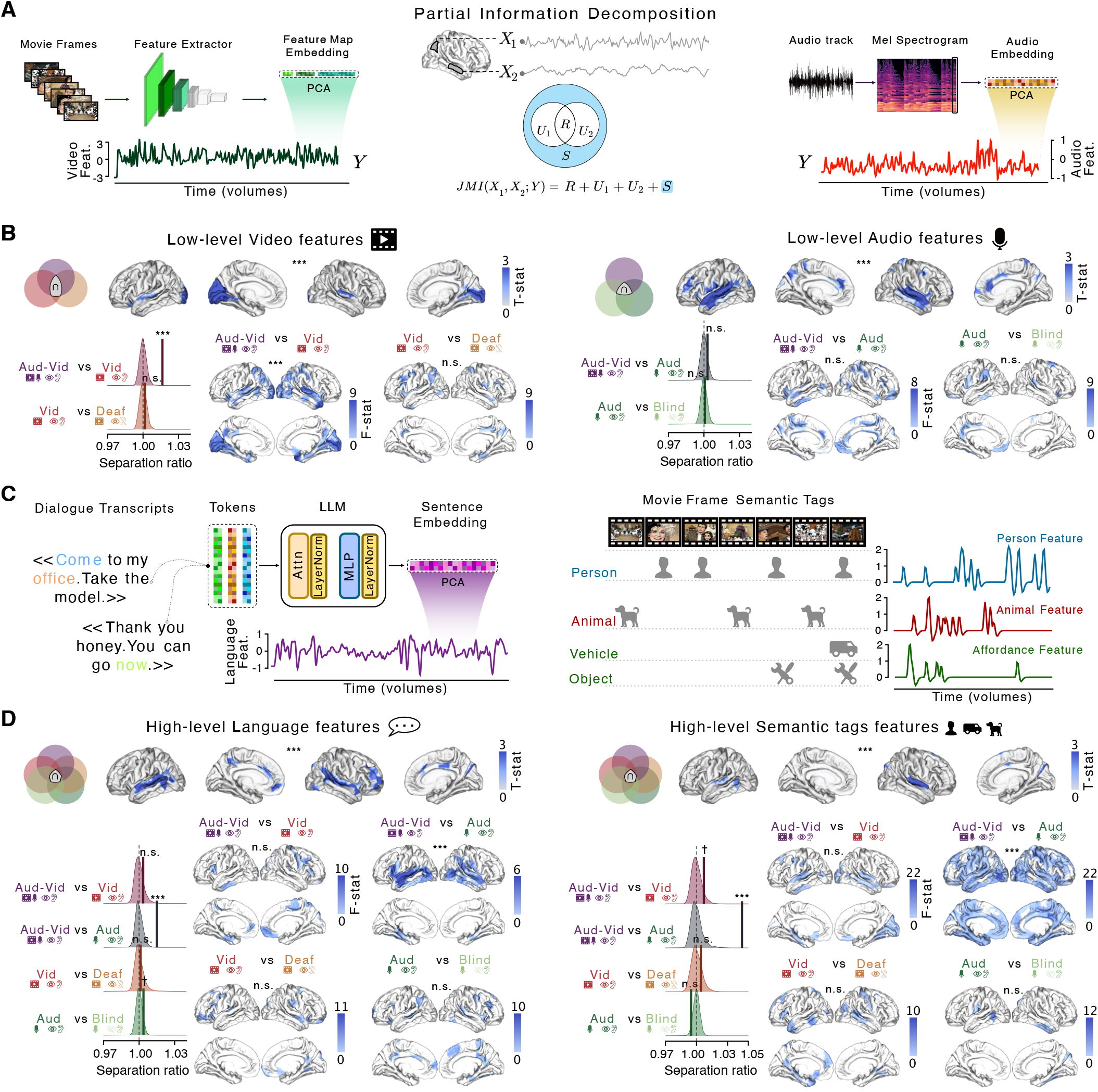
Partial Information Decomposition reveals synergistic encoding of low-level and high-level perceptual stimulus features. **A.** Schematic illustration of the Partial Information Decomposition (PID) analysis. For each pair of cortical regions, PID decomposed the joint mutual information between regional BOLD activity and stimulus features over the course of the narrative into informational atoms and here we showed the synergistic component. Low-level visual features were extracted from movie frames using a feature extractor deep visual model, whereas low-level auditory features were extracted from the audio track using Mel-spectrogram representations. **B**. Top, conjunction analyses identify low-level visual or audio encoding patterns common to the relevant experimental groups, revealing cortical regions whose synergistic encoding was invariant to sensory modality or sensory experience. Statistical significance was determined by cluster correction and permutation testing. Venn diagrams indicate the groups included in each conjunction analysis. Bottom, separation-ratio plots show observed contrast statistics against permutation null distributions among contrasted whole-brain synergistic interaction profiles. Brain maps show regions whose seed-to-whole-brain low-level feature-encoding interaction profiles differed between conditions, estimated using fcMVPA. Asterisks and transparency indicate statistical significance (P<0.05, cluster-corrected for brain plots). When fcMPVA showed no significant clusters, we highlighted regions with an F-statistic greater than 3. **C**. Schematic illustration of high-level feature extraction. Language features were extracted from dialogue transcripts using large language model (LLM) embeddings, while semantic-tag features were derived from frame-level annotations of high-level movie content, including person, animal, vehicle and object categories. **D**. Top, conjunction analyses identify high-level language or semantic tags encoding patterns common to all experimental groups. Bottom, separation-ratio plots and brain maps showing high-level feature-encoding interaction profiles differed between conditions. Legend: * P<0.05, ** P<0.01, *** P<0.001; n.s., not significant. Due to copyright, the original movie frames have been replaced with images generated using deep image generative models.

This approach determined whether stimulus information was duplicated across regions or instead emerged only through their joint activity, thus directly linking distributed neural interactions to the encoding of low-level and high-level stimulus content. Joint mutual information analyses revealed distributed cortical hubs that were broadly consistent with the nature of the encoded stimulus content (Fig. S3). Notably, although synergistic (Fig. S4) and redundant (Fig. S5) interactions showed highly distinct interaction topologies when normalized relative to JMI, their large-scale cortical hub organization was substantially more similar, revealing partially shared cortical architectures supporting complementary forms of distributed perceptual encoding. To characterize these interaction profiles, we used the same analytical framework previously applied to IID, including conjunction analyses to identify invariant encoding patterns across experimental groups, as well as separation-ratio and brain-wide connectome inference analyses to quantify condition-dependent differences at both whole-brain and regional scales.

We first focused on low-level perceptual features extracted from the frames and the audio track of the movie (Fig. 4A). Low-level visual features were extracted from early layers of a deep visual model trained for object recognition, whereas low-level auditory features were computed using Mel-spectrogram time-frequency representations. Conjunction analyses revealed invariant synergistic encoding patterns shared across the relevant experimental groups for both visual (all clusters P < 0.0062 cluster-corrected) and auditory (all clusters P < 0.0046 cluster-corrected) low-level features, indicating that distributed encoding of sensory information was partially preserved independently of sensory richness and previous sensory experience. By marginalizing the interactions for each brain area, we found that low-level visual features were consistently encoded mostly in occipital cortices, whereas low-level auditory features were consistently encoded primarily in temporal and frontal regions across the relevant groups (Fig. 4B).

Notably, similar invariant interaction patterns across relevant groups were also observed for redundant interactions (Fig. S6A top, all clusters P < 0.0092 cluster-corrected for both audio and visual low-level features). Comparable cortical organizations also emerged when conjunction analyses were applied directly to JMI (Fig. S7 top, all clusters P < 0.0014 cluster-corrected for both audio and visual low-level features), although low-level auditory feature encoding appeared spatially broader in this case. These results suggest that the large-scale organization of redundant and synergistic low-level feature encoding patterns may be partially constrained by the overall amount of stimulus-related low-level information across brain areas, despite redundancy and synergy being normalized relative to total joint mutual information.

Condition-dependent differences in low-level feature encoding were comparatively limited (Fig. 4B, bottom). For synergistic interactions, reliable whole-brain separation emerged only for low-level video features between audiovisual and visual-only conditions (SR = 1.02, P = 0.0002), with brain-wide connectome inference analyses localizing these effects primarily to occipital and temporal cortices (all clusters P < 0.002 cluster-corrected, all ηp^2^ > 0.56). Similar patterns were observed for redundant interactions (Fig. S6A bottom, SR = 1.02, P = 0.0002), although effects were relatively more pronounced within temporal regions (all clusters P < 0.008 cluster-corrected, all ηp^2^ > 0.52). All remaining low-level feature contrasts showed no significant differences across experimental groups in both synergistic and redundant interactions (all P > 0.05).

We next investigated high-level perceptual features capturing the semantic and linguistic structure of the narrative (Fig. 4C). High-level language features were extracted from the movie dialogue transcripts using sentence embeddings derived from a large language model (LLM), whereas high-level semantic-tag features were obtained from frame-level annotations describing the presence of semantic categories such as people, animals, vehicles and objects.

Conjunction analyses revealed invariant synergistic encoding patterns shared across all experimental groups for both linguistic (all clusters P < 0.0002 cluster-corrected) and semantic-tag features (all clusters P < 0.0002 cluster-corrected), indicating that distributed encoding of high-level information was largely preserved independently of current sensory modality and previous sensory experience (Fig. 4D top). Language features were synergistically encoded mostly in frontal and temporal regions, while semantic-tag features most prominently in temporal areas. Similar invariant encoding patterns were also observed for redundant interactions (Fig. S6B top, all clusters P < 0.0002 cluster-corrected for both high-level features) and joint mutual information (Fig. S7 bottom, all clusters P < 0.0002 cluster-corrected for both high-level features), again suggesting partially shared cortical architectures supporting complementary forms of high-level distributed perceptual encoding.

When examining the effects of sensory richness and sensory deprivation on high-level perceptual feature encoding (Fig. 4D, bottom), synergistic interactions showed reliable whole-brain separation only between audiovisual and auditory-only conditions for both language (SR = 1.01, P = 0.0004) and semantic-tag features (SR = 1.04, P = 0.0002). Brain-wide connectome inference analyses localized these effects primarily to temporal and frontal cortices for language features (all clusters P < 0.001 cluster-corrected, all ηp^2^ > 0.55) and to broadly distributed cortical regions spanning all cerebral lobes for semantic-tag features (all clusters P < 0.0002 cluster-corrected, all ηp^2^ > 0.60). Notably, no significant differences emerged between typically developed and sensory-deprived participants under matched unimodal stimulation (all P > 0.05).

Redundant interactions showed additional significant contrasts (Fig. S6B bottom). Beyond the audiovisual versus auditory-only comparison (SR > 1.01, P < 0.0006 for both high-level features), reliable whole-brain differences also emerged between typically developed and blind participants for language features during auditory-only stimulation (SR = 1.01, P = 0.028), and between audiovisual and visual-only conditions for semantic-tag features (SR = 1.01, P = 0.044). For the audiovisual versus auditory-only comparison, brain-wide connectome inference maps were broadly similar to those observed for synergistic interactions (all clusters P < 0.0002 cluster-corrected, ηp^2^ > 0.61 for both high-level features). By contrast, language-feature differences between auditory-only TD and blind participants (all clusters P < 0.005 cluster-corrected, ηp^2^ > 0.52) and semantic-tag differences between audiovisual and visual-only conditions (all clusters P < 0.037 cluster-corrected, ηp^2^ > 0.51) were localized primarily to occipito-temporal cortices. No significant effects were observed for language features in the audiovisual versus visual-only and visual-only versus deaf comparisons, nor for semantic-tag features in the sensory deprivation contrasts under matched unimodal stimulation (all P > 0.05).

Taken together, these results reveal that distributed neural interactions during naturalistic perception encode both low-level sensory features and high-level semantic information through complementary redundant and synergistic processes. Cortical hubs of information dynamics remained closely linked to modality-specific sensory systems for low-level feature encoding, whereas high-level features were encoded across associative transmodal areas. More broadly, these findings show how large-scale neural interactions support perception by distributed processing across the human brain, from sensory representations to high-level semantic content.

## Discussion

Perception in natural environments requires the processing of continuous sensory input to form coherent representations of the surrounding world. Rather than relying on isolated cortical modules or a simple serial progression from sensory to association cortices 70,71, this process likely emerges from distributed large-scale interactions spanning multiple brain areas 12,72–77.

Our results strongly support this view by showing that naturalistic vision and audition rely on complementary forms of distributed information processing, shaped by both current sensory input and lifelong sensory experience. In this context, our findings contribute to current debates 12,13 on whether perceptual and cognitive computations are localized or distributed 9–12, and align with recent brain-wide recording studies showing widespread distributed neural populations recruited in perceptual behavior 19,20.

Consistent with recent work in resting-state activity 47,78, we found that conventional functional connectivity primarily reflected redundant interactions, indicating that a substantial fraction of classical interregional coupling corresponds to information shared across cortical regions. In contrast, synergistic interactions remained largely invisible to conventional connectivity analyses, despite revealing widespread distributed processing across the cortex. These results highlight how information decomposition approaches can uncover informational structures of large-scale brain dynamics that are not accessible through classical measures of interregional covariation alone. Importantly, we extend these observations from resting-state activity to naturalistic auditory and visual perception across variations in sensory modality and sensory developmental history.

The spatial organization of redundancy and synergy further aligned with the cortical hierarchy from unimodal sensory systems to association transmodal cortex 8,47. Redundant interactions were strongest within sensory systems and closely tracked the structure of available sensory input, consistent with a role in robust perceptual representation. Synergistic interactions instead were preferentially expressed across associative transmodal cortices and were comparatively more preserved across sensory conditions.

Because synergy captures information that emerges only through the joint activity of multiple regions, it provides a particularly strong signature of distributed processing 48,49,51,52,59. In this context, our results suggest that associative transmodal cortices support perceptual processing through distributed integrative dynamics spanning large-scale brain networks. These findings resonate with the proposed “synergistic core” of the brain 47, suggesting that distributed integrative architectures centered on the associative transmodal cortex remain engaged during perceptual processing despite substantial changes in sensory input and sensory developmental history.

A particularly important finding was the role of the prefrontal cortex as a candidate modality-independent hub of synergistic dynamics. Its interaction profiles were expressed across experimental conditions and groups, independently of whether the narrative was experienced through audiovisual, visual-only, or auditory-only stimulation, and irrespectively of congenital sensory deprivation. This stability suggests that the prefrontal cortex supports forms of distributed integrative processing that are not tied to a specific sensory channel. Rather, its synergistic interactions may coordinate partial signals from large-scale cortical sites into high-level semantic representations of perceptual content, potentially supporting the attentional and integrative operations required for coherent naturalistic cognition 79,80.

High-order interaction analyses extended this picture by showing that distributed processing during naturalistic perception is not limited to pairwise relationships. O-information was introduced to quantify whether multivariate systems are dominated by redundancy-like or synergy-like dependencies, and later fMRI work showed that synergistic cortical subsets can be widespread, high-order, and largely invisible to pairwise functional connectivity analyses 60,64. Multimodal perception elicited the strongest high-order synergy, suggesting that the simultaneous availability of visual and auditory information promotes collective computations that depend on multiple regions considered together. This provides an information-theoretic account of why multimodal perception may be especially effective in naturalistic contexts: it does not merely activate more sensory cortex, but strengthens an interaction-dependent, brain-wide processing architecture.

The PID feature-encoding analyses linked these distributed information dynamics to the perceptual content of the movie. Low-level visual and auditory features were encoded in largely modality-specific cortical systems, consistent with the known organization of sensory hierarchies and with naturalistic encoding studies showing that visual and auditory features are represented along sensory cortical pathways. Importantly, these low-level encoding patterns were observed for both redundant and synergistic components, suggesting that sensory information is both shared across regions and integrated through joint activity. This supports the idea that early perceptual representations combine robustness with integrative distributed processing, allowing sensory information to remain available across multiple cortical sites while also contributing to interaction-dependent computations.

High-level language and semantic features showed a different organization. Their encoding involved broader associative cortices and was more preserved across sensory modalities and sensory histories, especially for synergistic interactions. This pattern is consistent with evidence that narrative comprehension, semantic processing and event understanding recruit associative cortical areas that abstract away from the specific sensory format of the input. In our data, this abstraction was not simply expressed as local feature encoding, but emerged through distributed interactions among cortical regions. Thus, synergy may provide a mechanism by which the brain integrates partial sensory, linguistic and semantic signals into coherent high-level representations of naturalistic content.

Collectively, the PID results complement the IID and HOI findings by showing that the brain-wide architecture of redundant and synergistic interactions is not only a dynamical property of neural activity, but plays a fundamental role in carrying and coordinating stimulus-relevant information 49,52,56,59. Furthermore, these findings reveal how the encoding of perceptual information across the human brain, from low-level sensory features to high-level semantic meaning, balances sensory robustness, supported by redundancy across areas, with distributed processing supported by joint interactions among multiple regions 1,9,18.

Congenital sensory deprivation specifically reduces or reshapes the high-order synergistic architecture. Under matched unimodal input, SD participants show weaker high-order synergy than TD participants. Importantly, this does not imply that sensory-deprived brains lack distributed processing. Rather, deprivation appears to alter the balance between preserved modality-independent computations and the broader synergistic architecture normally refined by lifelong multisensory experience. Recent evidence from congenitally blind and deaf individuals suggests that meaningful and content-specific representations can emerge independently of typical sensory experience, pointing to the existence of an intrinsic organizational scaffold (or proto-architecture) that precedes and constrains experience-dependent specialization 65,81,82. Within this framework, cross-modal plasticity may reflect not a complete functional reassignment of deprived cortices, but a structured reweighting of pre-existing large-scale interactions grounded in intrinsic cortical geometry and supramodal organization. Accordingly, cortical systems may preserve modality-independent representational formats that flexibly support the integration of perceptual and semantic information across different sensory channels. Our findings are consistent with this view, showing that synergistic interactions within associative transmodal and prefrontal cortices remain comparatively preserved across sensory modalities and developmental histories, while the magnitude and extent of high-order distributed interactions are modulated by lifelong sensory experience.

These findings also inform current accounts of sensory deprivation and cortical plasticity. Rather than inducing unrestricted large-scale reorganization, congenital blindness and deafness appear to reshape cortical dynamics within the constraints imposed by intrinsic developmental architecture and pre-existing multisensory connectivity 83. Multisensory influences are known to reach not only association cortices but also areas traditionally considered unimodal, challenging strictly serial models of cortical processing. In this context, our results suggest that sensory deprivation preserves important components of naturalistic perception, particularly those related to modality-compatible sensory processing and high-level semantic structure, while selectively modifying the large-scale synergistic coordination normally reinforced through continuous multimodal experience. Thus, the functional reorganization may preserve the functional logic of distributed perceptual systems without fully reproducing the richness and integration supported by lifelong access to multiple sensory modalities.

More broadly, the present findings suggest that the richness of distributed synergistic processing depends not only on the number of available sensory channels, but also on the computational structure of sensory information itself. Vision and audition differ substantially in their representational geometry and information density 7,14,84,85. Visual input provides high-dimensional spatial information that can be processed across multiple cortical hierarchies 86, whereas auditory processing relies more strongly on temporally sequential and compressed representations 87. These differences may help explain why visual and especially audiovisual conditions elicited broader and stronger synergistic interactions than auditory-only stimulation. In this framework, multimodal perception may enhance distributed synergy not simply because more sensory information is available, but because audiovisual signals provide complementary high-dimensional information streams that can be jointly integrated across associative cortical networks 2. Conversely, the comparatively sequential structure of auditory representations may impose stronger constraints on large-scale synergistic integration. This interpretation is consistent with recent evidence showing that the representational richness and hierarchical compatibility of substituting sensory inputs shape the extent and depth of cross-modal plasticity in sensory-deprived brains 65,88.

In summary, our findings reveal that natural vision and audition is supported by a brain-wide information architecture organized around complementary forms of distributed processing. Across sensory modalities and sensory developmental histories, redundant interactions supported robust representations of perceptual information, whereas synergistic interactions coordinated distributed integration across large-scale cortical systems. By linking large-scale neural dynamics to the informational structure and perceptual content of cortical interactions, our results provide a framework for understanding how the human brain orchestrates the perceptual processing of continuous sensory to form coherent perceptual experience.

## Methods

### Participants

Fifty participants were recruited, including typically developed (TD) individuals and subjects with congenital sensory deprivation (SD). TD participants were assigned to one of three conditions: audiovisual (AV; *n* = 10, 35 ± 13 years, 8 females), auditory-only (A; *n* = 10, 39 ± 17 years, 7 females), or visual-only (V; *n* = 10, 37 ± 15 years, 5 females). The SD sample included congenitally blind participants presented with the auditory-only version (*n* = 11; 2 excluded for excessive head motion, final *n* = 9, 44 ± 14 years, 3 females) and congenitally deaf participants exposed to the visual-only version (*n* = 9, 24 ± 4 years, 5 females). All participants were right-handed, native Italian speakers, and had no history of neurological or psychiatric disorders. Deaf participants were proficient in Italian Sign Language and were not using hearing aids during testing; TD participants reported no hearing impairment, normal or corrected-to-normal vision, and no knowledge of Italian Sign Language. Additional information about the demographic data of the participants involved in the study can be found in recently published works ^65,66^.

### Stimulation and experimental procedure

Naturalistic stimulation consisted of edited visual, auditory, and audiovisual versions of the live-action movie *101 Dalmatians*. The movie was selected on the basis of several criteria. Its linear narrative structure and the predominance of dialogue-driven plot progression ensured that story comprehension remained intact across unimodal (visual-only or auditory-only) presentations. Crucially, the film depicts familiar social and physical environments such as human characters, domestic animals, indoor and outdoor settings, whose semantic content is accessible through multiple sensory modalities and consistent with the sociocultural context of the participant sample. This was particularly important given the inclusion of congenitally blind and deaf participants, for whom the stimulus needed to be semantically and experientially meaningful despite the absence of visual or auditory experience since birth. The movie also provides continuous, naturalistic variation in low-level sensory features, social content, and narrative context across its duration, enabling robust sampling of distributed brain dynamics. It was shortened to about 54 minutes and divided into six runs of approximately 8 minutes each, with 6-s fade-in and fade-out periods at the beginning and end of each run. To ensure equivalent narrative comprehension across participant groups with different sensory histories, each stimulus version was designed as a self-contained, modality-complete experience. In the auditory-only condition, a professional Italian audio description was superimposed on the original soundtrack for all participants, providing a verbal mirror of the visual content. In the visual-only and audiovisual conditions, Italian subtitles were displayed for all participants, providing a textual mirror of the auditory-linguistic content. This symmetric design ensured that none of the groups relied on sensory compensation absent from others within the same condition, and that narrative comprehension was matched across groups without introducing asymmetric confounds. A red fixation cross was displayed at screen center in all conditions involving visual presentation to standardize gaze behavior. Audio and video were delivered through MR-compatible headphones and LCD goggles. Sound mixing was performed with the LogicPro (Apple Inc., version 10.4) software. The video and audio clips were edited with iMovie (Apple Inc., version 10.1.10) software whereas for the creation of subtitles, we rely on the open-source cross-platform Aegisub 3.2.2 (http://www.aegisub.org/). Stimulation was administered through software package Presentation (Neurobehavioral System, Berkeley, CA, USA; version 16.5; http://www.neurobs.com).

Moreover, before scanning, participants rated their familiarity with the movie plot on a 1-to-5 Likert scale. During fMRI, each participant watched only one edited version of the movie (V, A, or AV) and was instructed to simply enjoy it. Structural and functional images were acquired in a single session. After scanning, participants completed an ad hoc two-alternative forced-choice questionnaire assessing comprehension of the story content; additional psychometric measures were also collected. Additional information about the stimulation and experimental design can be found in recently published works ^65,66^.

### fMRI data acquisition and preprocessing

Brain activity was recorded with a Philips 3T Ingenia scanner equipped with a 32-channel head coil. Functional images were acquired using gradient recall echo planar imaging (GRE-EPI; repetition time (TR) 2,000 ms; echo time (TE) 30 ms; flip angle (FA) 75°; field of view (FOV) 240 mm; acquisition matrix (in plane resolution) 80 × 80; acquisition slice thickness 3 mm; acquisition voxel size 3 × 3 × 3 mm; reconstruction voxel size 3 × 3 × 3 mm; 38 sequential axial ascending slices; total volumes 1,614 for the six runs of the movie, plus 256 for the control run). In the same session, a high-resolution anatomical scan was also collected using a T1-weighted magnetization-prepared rapid gradient echo sequence (TR 7 ms; TE 3.2 ms; FA 9°; FOV 224, acquisition matrix 224 × 224; slice thickness 1 mm; voxel size 1 × 1 × 1 mm; 156 sagittal slices).

fMRI data preprocessing was performed following the standard steps with AFNI (version 17.1.12) software package ^89^. First, we removed scanner-related noise correcting the data by spike removal (3dDe-spike). Then, all volumes comprising a run were temporally aligned and successively corrected for head motion using as base the first run (3dvolreg). A spatial smoothing with a Gaussian kernel (6 mm, full width at half maximum) was applied and then data of each run underwent percentage normalization. In addition, detrending applying Savitzky–Golay filtering (polynomial order: 3, frame length: 200 timepoints) was performed onto the normalized runs to smooth the corresponding time series and clean them from unwanted trends and outliers. Runs were then concatenated, and multiple regression analysis was performed to remove signals related to head motion parameters and movement spike regressors (framewise displacement above 0.3). Afterwards, single-subject fMRI volumes were non-linearly registered to the MNI-192 standard space ^90^. Cortical parcellation was performed using the HCP Atlas ^91^.

### Information-Theoretic Framework

Information theory provides a common language for quantifying statistical dependence among neural signals and between neural signals and external variables. In this framework, dependence is expressed as a reduction in uncertainty: one variable is informative about another when observing it makes the other less uncertain.

For a variable *Y*, Shannon entropy *H*(*Y*) quantifies uncertainty ^92^. Mutual information (MI) quantifies how much uncertainty about one variable or variable set is reduced after observing another. For two variables or variable sets *A* and *B*,

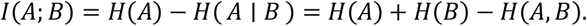

MI therefore gives a general measure of dependence that can be applied to individual regions, sets of regions, temporal states, or stimulus-derived variables. However, the total amount of information shared between variables does not by itself specify how that information is organized. The same total dependence may reflect information carried separately by individual variables, information shared across variables, or information that becomes available only when variables are considered jointly. The following sections introduce the specific decompositions used to distinguish these forms of information in the present analyses.

All information-theoretic quantities were estimated using Gaussian-copula mutual information (gcMI) ^67^. This estimator is well suited to continuous BOLD and stimulus-feature time series because it reduces sensitivity to non-Gaussian marginal distributions while preserving the empirical dependence structure among variables. For each variable *Y*, observations were rank-normalized through the empirical cumulative distribution function 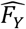 and mapped to standard-normal scores,

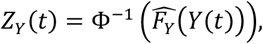

where Φ^−1^ is the inverse cumulative distribution function of the standard normal distribution. This transformation places all variables on a common Gaussian marginal scale while retaining their copula, that is, their multivariate dependence structure.

After copula normalization, information quantities were computed from covariance matrices. For a *d*-dimensional copula-normalized variable set *Z* with covariance matrix Σ_*z*_, Gaussian entropy is

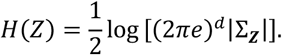

Accordingly, the mutual information between two copula-normalized variable sets *A* and *B* was evaluated as

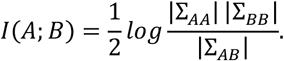

Here, Σ_*AA*_ and Σ_*BB*_ are the covariance matrices of the two variable sets, and Σ_*AB*_ is their full joint covariance matrix. Thus, all subsequent analyses are grounded in the same estimation principle: dependencies are quantified from the covariance structure of copula-normalized BOLD or stimulus-feature variables. The next sections specify how this common information-theoretic estimator was used to characterize temporal dependence, high-order multiregional structure, and stimulus-related information.

### Functional Connectivity

As a conventional reference for large-scale coupling, pairwise functional connectivity was computed between all cortical regions using Gaussian-copula mutual information. Each matrix entry quantified the statistical dependence between two regional BOLD time series sampled at the same movie time points.

For each participant and condition, the resulting symmetric FC matrix was converted into a whole-brain connectivity profile by vectorizing its upper triangular entries. Let ***p***_***a***_ and ***p***_***b***_ denote the connectivity profiles of two participants. Their dissimilarity was quantified using cosine distance This produced a participant-by-participant distance matrix for each contrast. The distance matrix was projected into two dimensions using multidimensional scaling (MDS) for visualization only; statistical testing was performed on the original cosine-distance structure.

For each contrast, group separation was quantified using a separation ratio. Let *G*_1_ and *G*_2_ denote the two groups being compared and let *y*_*a*_ indicate the group label of participant *a*. The mean between-group distance was computed across all participant pairs belonging to different groups,

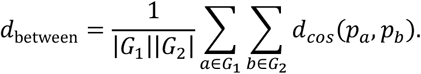

The mean within-group distance was computed across all unique participant pairs belonging to the same group,

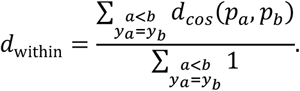

The separation ratio was then defined as SR = *d*_between_ / *d*_within_. Values above one indicate that participants from the two contrasted groups had more distinct whole-brain FC profiles than expected from within-group variability. Statistical significance was assessed by permutation testing, in which group labels were shuffled to generate a null distribution of SR values for each contrast.

The same profile-analysis procedure was subsequently applied to interaction-based information decomposition subsequent analyses. In particular, across all matrix-based analyses, MDS and SR refer to the same steps: vectorization of participant-level matrices, cosine-distance computation between participant profiles, MDS visualization, and permutation testing of SR.

For circular graph visualizations, the strongest connections were displayed for each group-average matrix. Brain maps were computed from the corresponding weighted networks using betweenness centrality, providing a compact visualization of cortical regions occupying central positions within the interaction profile.

### Integrated Information Decomposition

Integrated Information Decomposition (IID) was used to decompose temporally persistent information in the BOLD dynamics of pairs of cortical regions ^47,53^. Whereas functional connectivity measures equal-time dependence between two regional time series, IID asks how the past state of a two-region BOLD system constrains its future state, and whether this temporal information is redundant or synergistic.

For each participant, condition, and pair of cortical regions (*X*_*i*_, *X*_*j*_), we computed the lag-1 time-delayed mutual information (TDMI),

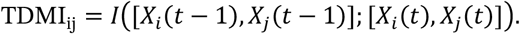

TDMI quantifies how much information the previous joint state of the two regions carries about their next joint state. It was estimated from the Gaussian-copula covariance of the past-future BOLD bivariate random variable. IID decomposes TDMI into informational nodes defined over both the past and the future. Let ℳ = {*R, U*_*i*_, *U*_*j*_, *S*} denote the four bivariate information nodes: redundancy, information unique to region *i*, information unique to region *j*, and synergy. IID defines atoms 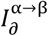 for α, β ∈ ℳ, where α denotes the informational mode in the past and β denotes the informational mode in the future. The TDMI is recovered by summing the 16 atoms: 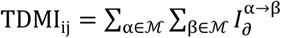.

Because the full 16-atom system is not determined by a single-target redundancy function alone, IID requires an additional double-redundancy constraint. Following the Gaussian minimum-mutual-information formulation used in previous fMRI applications, the redundancy-to-redundancy atom was defined as the minimum mutual information across the four pairwise mutual information linking each past variable to each future variable,

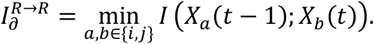

This term quantifies information that is redundantly available across the two regions in the past and remains redundantly available across the two regions in the future. The remaining IID atoms were obtained by solving the linear system imposed by the product-lattice consistency constraints.

From the full decomposition, we retained the two diagonal persistence atoms, 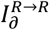 and 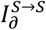. Persistent redundancy, 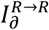, quantifies future BOLD information that is shared across the two regions and remains shared across successive time points. Persistent synergy, 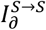, quantifies future BOLD information that is available only from the joint state of the two regions and remains synergistically represented over time. By focusing on these diagonal atoms, the analysis isolated temporally stable redundant and synergistic components, rather than information that was copied, transferred, or transformed between informational nodes.

The persistence atoms were estimated separately for every participant, condition, and ROI pair, yielding subject-level symmetric redundancy and synergy matrices *R*^per^ and *S*^per^. The resulting matrices were entered into the matrix-profile, conjunction, and fcMVPA procedures described in Statistical analysis section. Furthermore, we also computed a redundancy-to-synergy interaction gradient: edge weights were rank-normalized within each information matrix to make redundancy and synergy comparable in scale, and a signed edge-level contrast was computed as

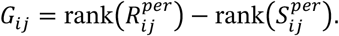

Positive values indicate redundancy-dominated interactions, whereas negative values indicate synergy-dominated interactions. The interaction gradient was used to visualize the cortical distribution of redundancy and synergy within IID, as well as for the conjunction analysis (see Statistical Analysis for details).

### High-order interactions

Pairwise IID quantifies temporally persistent redundancy and synergy between two regions. However, naturalistic perception may also involve larger multiregional ensembles in which information is not reducible to pairwise interactions. To test whether synergistic structure extended beyond pairs into larger brain subsystems, we quantified high-order interactions using O-information^60^. For a subset *B* of *k* cortical regions, let 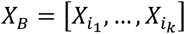 denote the corresponding multiregional BOLD subsystem. O-information is defined as the difference between total correlation and dual total correlation,

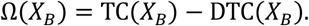

Total correlation measures the overall dependence among the regions in the subset relative to a model in which all regional time series are independent,

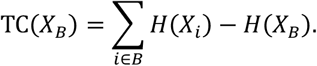

Dual total correlation measures dependence from the complementary perspective of how much uncertainty remains in each regional signal after conditioning on the rest of the subsystem,

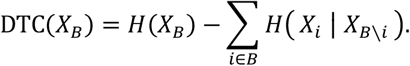

O-information compares these two multivariate dependence structures. Positive values indicate redundancy-dominated high-order structure, whereas negative values indicate synergy-dominated high-order structure. Thus, in the present analysis, more negative Ω values correspond to stronger high-order synergy across the selected cortical subsystem. Under the Gaussian-copula model, the entropy terms entering TC and DTC were evaluated from the covariance matrix of the copula-normalized regional BOLD signals in each subset. This allowed O-information to be computed for candidate multiregional subsystems using the same covariance-based estimator as the other information-theoretic analyses.

For each experimental condition, high-order interactions were estimated from the group-level covariance structure obtained by averaging participant-level covariance matrices. Because the number of possible subsets grows combinatorially *k*, exhaustive search over all region combinations was not feasible. We therefore used simulated annealing ^64^ to identify maximally synergistic subsystems, defined as subsets minimizing Ω(*X*_*B*_).

The search was performed separately for each condition and target subset size *k* ∈ {3,6,8,10,12,14,16,18,20,22}. For each condition and subset size, 200 independent simulated-annealing runs were initialized from random subsets of size *k*, each implementing a maximum of 10 000 optimization iterations. At each iteration, a candidate subset was generated by replacing one region in the current subset with one region outside the subset. Moves that decreased O-information, and therefore increased high-order synergy, were accepted deterministically. Moves that increased O-information could still be accepted with a probability controlled by the annealing temperature, allowing the algorithm to escape local minima early in the search and become increasingly selective as the temperature decreased. Each run terminated when convergence criteria were met or when no further beneficial move was available.

The 200 optimized solutions for each condition and subset size were interpreted as independent explorations of the same group-level synergy landscape, not as independent sample observations. For each solution, the final O-information value was retained. The distribution of optimized values across subset sizes defined the high-order synergy landscape of each experimental condition, and the optimal interaction order was defined as the subset size yielding the most negative mean O-information.

To characterize the anatomical composition of maximally synergistic subsystems, we computed the participation frequency of each cortical region across optimized solutions. Participation maps were weighted by synergy strength, using −Ω for synergy-dominated subsets, so that regions recurring in more strongly synergistic solutions contributed more strongly to the final map. Group comparisons were performed across the O-information landscapes, and optimal interaction orders were defined as the subset sizes yielding maximal synergy, i.e., the most negative mean O-information, within each condition.

### Partial Information Decomposition

Partial Information Decomposition (PID) was used to determine what stimulus information was encoded redundantly or synergistically by pairs of brain regions ^57–59^. In contrast to IID, which decomposes intrinsic temporal dependencies in BOLD dynamics, PID decomposes the information that two neural sources provide about an external movie-feature target.

For each participant, condition, ROI pair (*X*_*i*_, *X*_*j*_), and stimulus-feature target *T*, we first estimated the joint mutual information between the two regional BOLD time series and the feature,

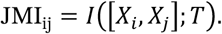

This quantity measures how much information about the feature target is carried by the joint state of the two regions. PID decomposes this joint information into redundant, unique, and synergistic components,

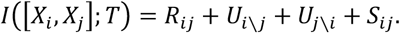

Here, *R*_*ij*_ is the information about the feature that is redundantly available from both regions, *U*_*i*\*j*_ and *U*_*j*\*i*_ are the feature information uniquely provided by each region, and *S*_*ij*_ is the feature information available only when the two regional time series are considered jointly.

We used the Gaussian minimum-mutual-information PID. Redundancy was defined as

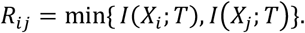

The unique terms were *U*_*i*\*j*_ = *I*(*X*_*i*_; *T*) − *R*_*ij*_(*T*), and *U*_*j*\*i*_ = *I*(*X*_*j*_; *T*) − *R*_*ij*_(*T*). Synergy can then be derived as

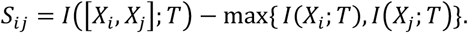

Thus, redundancy quantifies overlapping stimulus information carried by both regions, whereas synergy quantifies stimulus information that cannot be recovered from either region alone but becomes available when both regions are analyzed together.

PID was applied separately to every ROI pair and feature target, yielding participant-level symmetric matrices of joint mutual information, redundancy, and synergy. The resulting matrices were entered into the matrix-profile, conjunction, and fcMVPA procedures described in Statistical analysis.

### Stimulus-feature Target Construction

Stimulus-feature targets were constructed to quantify how distributed neural interactions encoded low-level sensory information and high-level perceptual content in the movie. All features were extracted at their native temporal resolution and then temporally aligned to the fMRI signal using the same preprocessing logic. When features were sampled more densely than the BOLD time series, they were first downsampled to the fMRI temporal resolution and then convolved with a canonical hemodynamic response function to account for the delayed BOLD response. For high-dimensional feature spaces, dimensionality reduction was applied using principal component analysis, and the first principal component was retained as a scalar target time course for PID analysis.

High-level language features were derived from the movie dialogue transcript using ChatGPT ^93^ embeddings. Text segments were passed through the language encoder to obtain semantic embedding vectors at each corresponding timestamp. These embeddings represented the evolving linguistic and semantic content of the narrative, including word meaning, sentence context, and discourse-level information. After temporal alignment with the BOLD signal and hemodynamic convolution, the embedding space was reduced with principal component analysis, yielding a single language-feature target time course.

High-level semantic-category features were obtained from event annotations marking the presence of broad content categories in the movie. Binary boxcar regressors were created for time periods in which persons, animals, vehicles, or objects appeared. Vehicle and object annotations were merged into a single affordance-related category, reflecting visually actionable or manipulable entities in the scene. This produced three semantic event regressors: person, animal, and affordance. Each regressor was downsampled, hemodynamically aligned to the BOLD signal, and the resulting time courses were stacked to form a multivariate semantic-category target.

Low-level visual features were extracted from movie frames using a pretrained VGG-19 network ^94^. Each frame was passed through the visual encoder, and activations from the first three visual-processing layers were used to describe frame-wise image properties. These activations captured low-level visual structure such as local edges, textures, contrast patterns, and spatial configurations. The resulting frame-wise embedding sequence was aligned to the BOLD time series and reduced with principal component analysis, yielding a single visual-feature target time course.

Low-level auditory features were extracted from the complete movie audio track using a Mel-spectrogram representation. This representation captured the time-varying spectral energy of the soundtrack on a perceptually motivated frequency scale, providing a compact description of acoustic properties such as intensity, frequency content, and temporal modulation. The Mel-spectrogram was temporally aligned to the fMRI signal, convolved with the hemodynamic response function, and reduced with principal component analysis to obtain a single auditory-feature target time course.

### Statistical analysis

The principal group contrasts were defined to isolate two factors: current sensory richness and lifelong sensory experience. Sensory-richness contrasts compared typically developed audiovisual participants with typically developed visual-only participants, and typically developed audiovisual participants with typically developed auditory-only participants. Sensory-experience contrasts compared typically developed visual-only participants with congenitally deaf participants under matched visual stimulation, and typically developed auditory-only participants with congenitally blind participants under matched auditory stimulation. Unless otherwise specified, all statistical tests were performed independently for each contrast and analysis family.

For all matrix-based analyses, participant-level matrices were converted into whole-brain profiles by vectorizing their upper triangular entries. This common procedure was applied to functional connectivity matrices, IID-derived persistent redundancy and persistent synergy matrices, and PID-derived joint mutual information, redundancy, and synergy matrices.

For brain-wise localization of condition-dependent effects, we used functional connectivity multivariate pattern analysis (fcMVPA) ^95^. This procedure was applied to the same matrix families described above. For each seed region, the vector of its connections to all other cortical regions defined a seed-specific whole-brain interaction fingerprint. For each contrast, participant-level fingerprints from the two groups were concatenated into a subject-by-edge matrix. Singular value decomposition was applied to this matrix, and the top 5 left-singular vectors were retained as a low-dimensional representation of each participant’s seed-specific interaction profile. Between-group differences were tested on these components using the Wilks-derived *F* statistic. The analysis was repeated independently for each seed region and contrast, yielding brain-wise statistical maps that were corrected using cluster-based permutation testing. Cluster formation used a cluster-alpha threshold of 0.05, and clusters were retained when their permutation-corrected p-value was below 0.05. For significant clusters, effect size was quantified as the mean partial eta-squared across ROIs in the cluster,

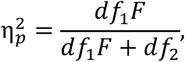

where *df*_1_ and *df*_2_ are the numerator and denominator degrees of freedom of the Wilks-derived *F* statistic.

Conjunction analyses were performed using a direct cluster-based permutation framework to identify interaction or feature-encoding patterns jointly preserved across experimental groups ^96,97^. For each group, subject-level maps were converted into one-sample t-statistic maps, and conjunction statistics were computed by retaining, at each cortical region, the minimum t-value shared across groups for positive effects (or the maximum shared negative t-value for negative effects). Cortical regions exceeding the conjunction threshold were grouped into spatially contiguous clusters based on the adjacency matrix. Cluster significance was assessed using nonparametric permutation testing with random sign-flipping at the subject level within each group. For each null permutation, conjunction statistics and cluster-level statistics were recomputed, and the maximum cluster statistic across the cortex was retained to form a null distribution controlling the family-wise error rate. Observed conjunction clusters were considered significant if their cluster-level statistic exceeded the permutation-derived null distribution threshold of 0.05.

This procedure therefore tested for cortical effects that were simultaneously significant across all groups included in the conjunction while accounting for spatial dependence and multiple comparisons within a unified permutation framework. For IID, conjunction analyses were applied to the redundancy-to-synergy interaction gradient to identify cortical hubs whose position along the redundancy–synergy axis was shared across experimental groups. For PID, conjunction analyses were applied separately to joint mutual information, redundancy, and synergy for each stimulus-feature family. Low-level sensory conjunctions included only groups for which the corresponding sensory stream was available, whereas high-level language and semantic-tag conjunctions included all experimental groups.

Conjunction analyses were also computed using a more conventional intersection method to test the robustness of the findings in the IID results ^96,97^. For each group included in a conjunction, the relevant subject-level maps were converted into statistical maps and corrected using cluster-based permutation testing. Corrected significance maps were then intersected across groups, and a cortical region was retained only if it survived the corrected threshold in every group included in the conjunction.

For high-order interaction analyses, statistical inference was performed on the one-dimensional O-information landscapes obtained across ROI subset sizes. We performed permutation cluster-based testing to identify consecutive ROI subset sizes at which conditions differed in maximal achievable synergy. Four contrasts were assessed independently: audiovisual TD participants versus video-only TD participants, audiovisual TD participants versus audio-only TD participants, video-only TD participants versus congenitally deaf participants, and audio-only TD participants versus congenitally blind participants. Cluster formation used a cluster-alpha threshold of 0.05, and clusters were retained when their permutation-corrected p-value was below 0.05. For significant one-dimensional clusters, effect size was computed as the mean Cohen’s *d* across all subset sizes included in the cluster. In addition, condition differences at the optimal interaction order were assessed by comparing O-information values across independent optimization runs at the subset size yielding maximal mean synergy.

To assess the relationship between conventional functional connectivity and IID-derived interactions, participant-level functional connectivity profiles were correlated with the corresponding persistent redundancy and persistent synergy profiles using vectorized upper-triangular matrix entries. Correlations were computed separately for each participant and experimental group, and group-level significance was assessed against the null hypothesis that profile correlations did not differ from zero.

Across all cluster-based analyses ^98,99^, corrected significance was defined as permutation-corrected (p < 0.05). For profile-level analyses, significance was assessed using permutation null distributions of the separation ratio. For brain-wise fcMVPA clusters, reported effect sizes correspond to the mean partial eta-squared across all ROIs in the cluster. For one-dimensional high-order interaction clusters, reported effect sizes correspond to the mean Cohen’s *d* across all subset sizes in the cluster.

## Acknowledgements

This research received funding from the European Research Council under the Grant No. 820213 (ThinkAhead), the Italian National Recovery and Resilience Plan (NRRP), M4C2, funded by the European Union, NextGenerationEU (Project IR0000011, CUP B51E22000150006, “EBRAINS-Italy”), and the Ministry of University and Research, PRIN PNRR P20224FESY and PRIN 20229Z7M8N. The funders had no role in study design, data collection and analysis, decision to publish, or preparation of the manuscript. We used a Generative AI model to correct typographical errors and edit language for clarity.

## Data and code availability

All fMRI data are available at https://figshare.com/s/de0d52bf08280cf0c4bd ^66^. We used Nilearn for fMRI analysis, which is available at https://nilearn.github.io/. We also used the GCMI library openly available at https://github.com/robince/gcmi. Moreover, we used the phyid library for the computation of the Integrated Information Decomposition analysis, which is openly available at https://github.com/Imperial-MINDlab/integrated-info-decomp.

## Author contributions

*: conceptualization, software, methodology, formal analysis, validation, data curation, visualization, writing - original draft, writing - review and editing.

*: conceptualization, software, methodology, formal analysis, validation, data curation, visualization, writing - original draft, writing - review and editing.

conceptualization, data curation, writing - review and editing.

^†^: conceptualization, data curation, writing - review and editing.

^†^: conceptualization, funding acquisition, methodology, project administration, supervision, validation, writing - original draft, writing - review and editing.

## Competing interests

The authors declare no conflict of interests.

## Supplementary Materials

**Figure S1.**
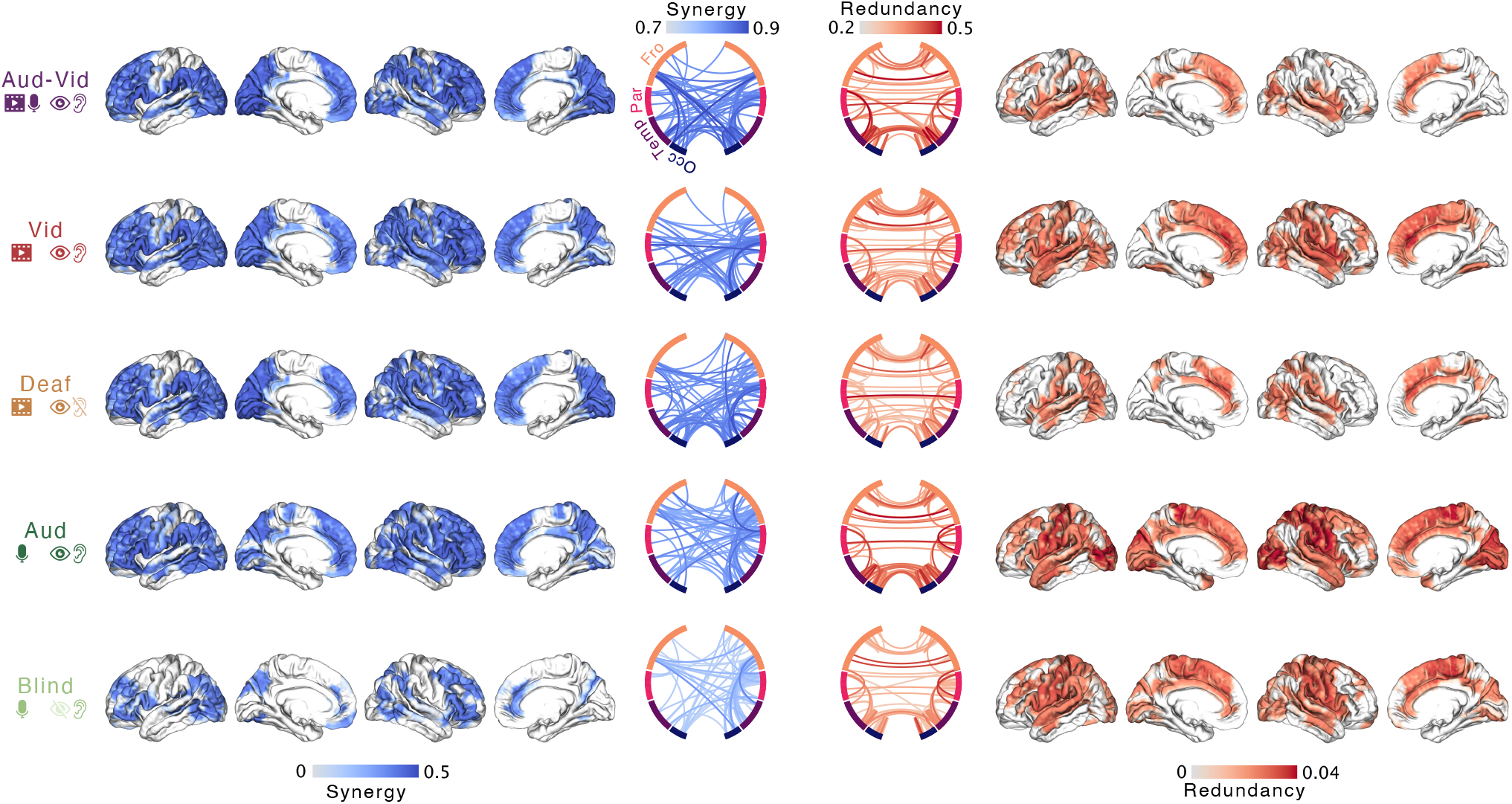
Brain-wide distribution of persistent synergy and redundancy across experimental groups. Group-average temporally persistent synergy and redundancy atoms estimated with Integrated Information Decomposition (IID). Circular plots showing redundancy and synergy scores for the top 100 interactions, while brain maps show regional betweenness centrality of each cortical region. These informational values provide the full condition-wise distribution of redundant and synergistic temporal interactions underlying the redundancy-to-synergy gradient.

**Figure S2.**
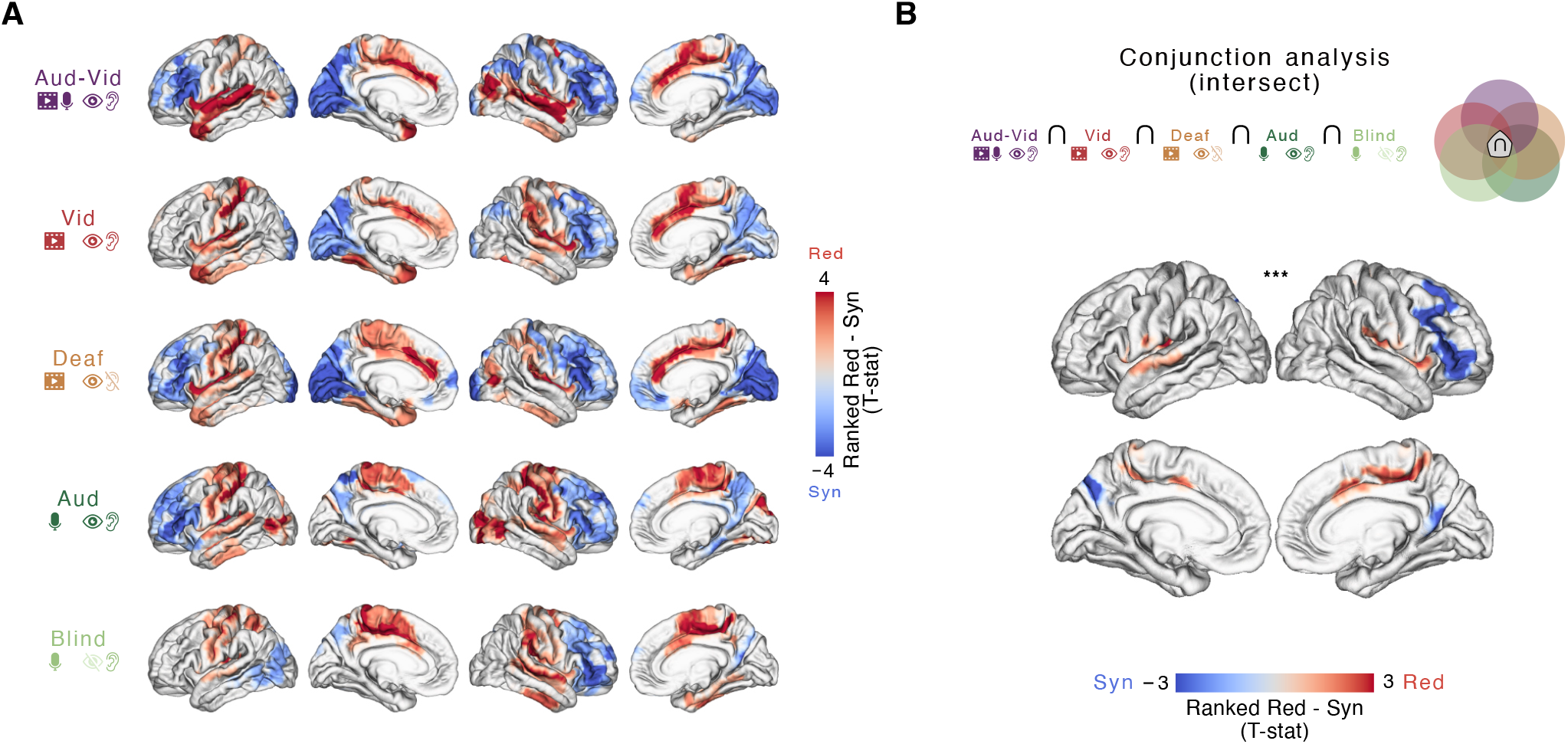
Redundancy-to-synergy gradients across experimental groups and intersect conjunction analysis. **A**. Brain maps showing significant ranked redundancy-minus-synergy gradients for each experimental group, expressed as t-statistics across cortical regions. Positive values indicate redundancy-dominated regions, whereas negative values indicate synergy-dominated regions. Transparency indicates statistical significance. **B**. Conjunction analysis identifying redundancy-to-synergy gradients shared across all experimental groups using a p-value intersection criterion. For each group, cluster correction was first applied separately to the corresponding t-maps, and the corrected p-values were then intersected across groups to identify cortical regions surviving the conjunction criterion (P<0.05). Brain maps show the resulting conjunction t-statistics, with asterisks and transparency indicating statistical significance.

**Figure S3.**
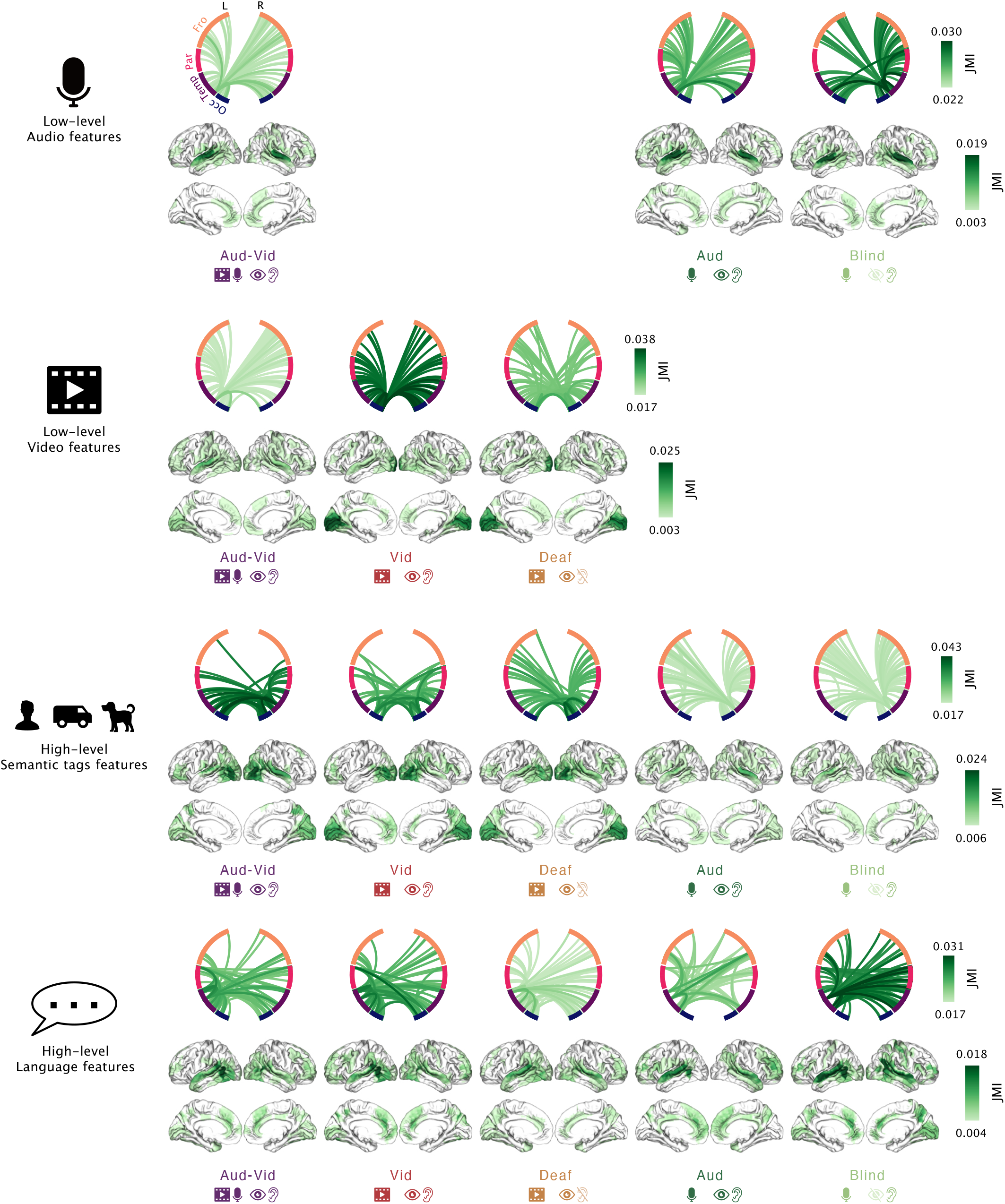
Joint mutual information between brain interactions and stimulus features across experimental groups. Group-average joint mutual information (JMI) between pairs of cortical regions and stimulus features, estimated for each experimental group and feature class. Circular plots show the top 100 feature-encoding interactions, while brain maps show regional betweenness centrality of each cortical region based on the corresponding JMI interaction profile. Rows correspond to low-level audio features, low-level video features, high-level semantic-tag features and high-level language features. These maps provide the full condition-wise distribution of joint feature information underlying the Partial Information Decomposition analyses.

**Figure S4.**
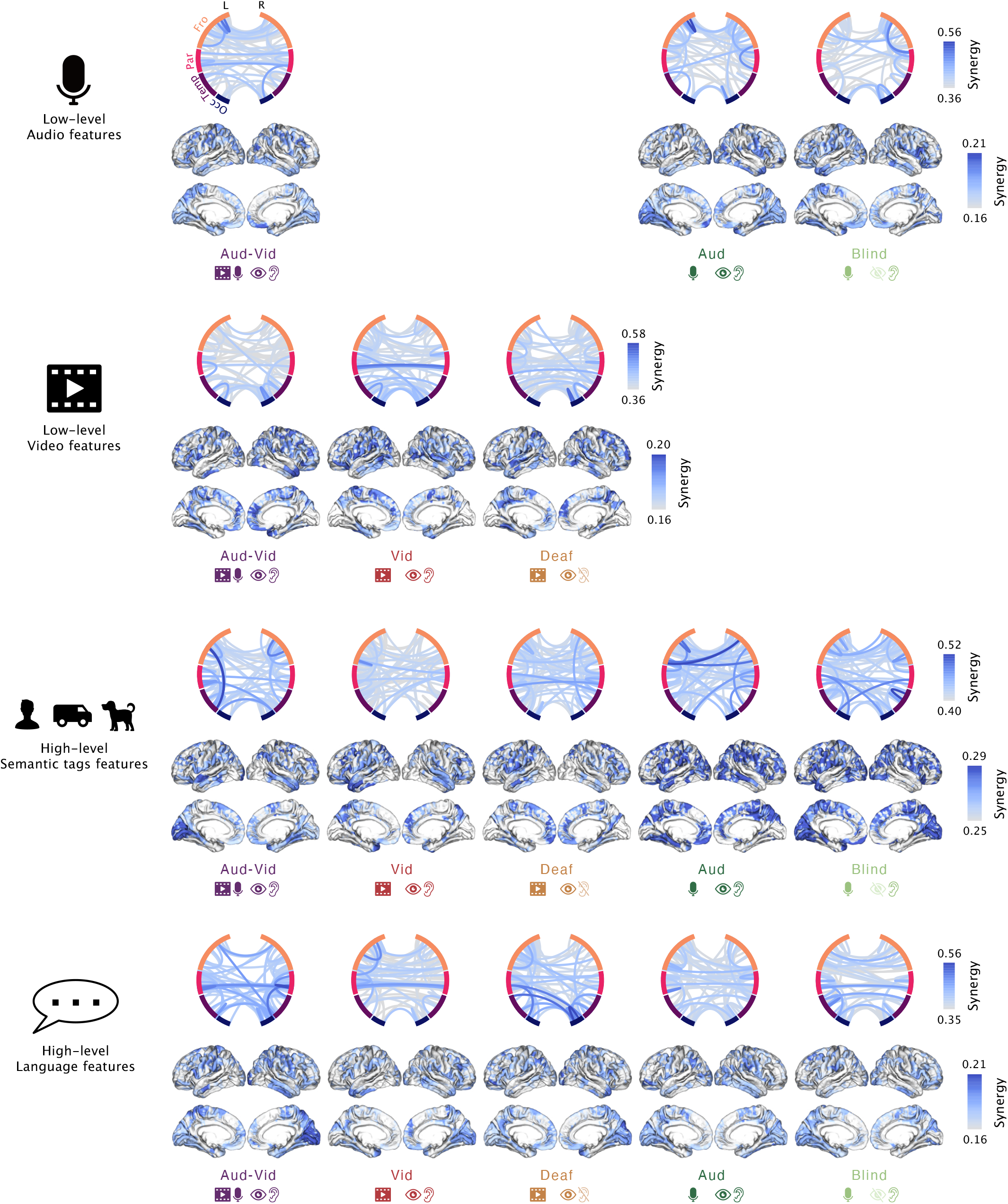
Synergistic feature encoding across experimental groups and stimulus features. Group-average synergistic information between pairs of cortical regions and stimulus features, estimated with Partial Information Decomposition (PID) for each experimental group and feature class. Circular plots show the top 100 synergistic feature-encoding interactions, while brain maps show regional betweenness centrality of each cortical region based on the corresponding synergy interaction profile. Rows correspond to low-level audio features, low-level video features, high-level semantic-tag features and high-level language features. These maps provide the full condition-wise distribution of synergistic feature encoding underlying the PID analyses.

**Figure S5.**
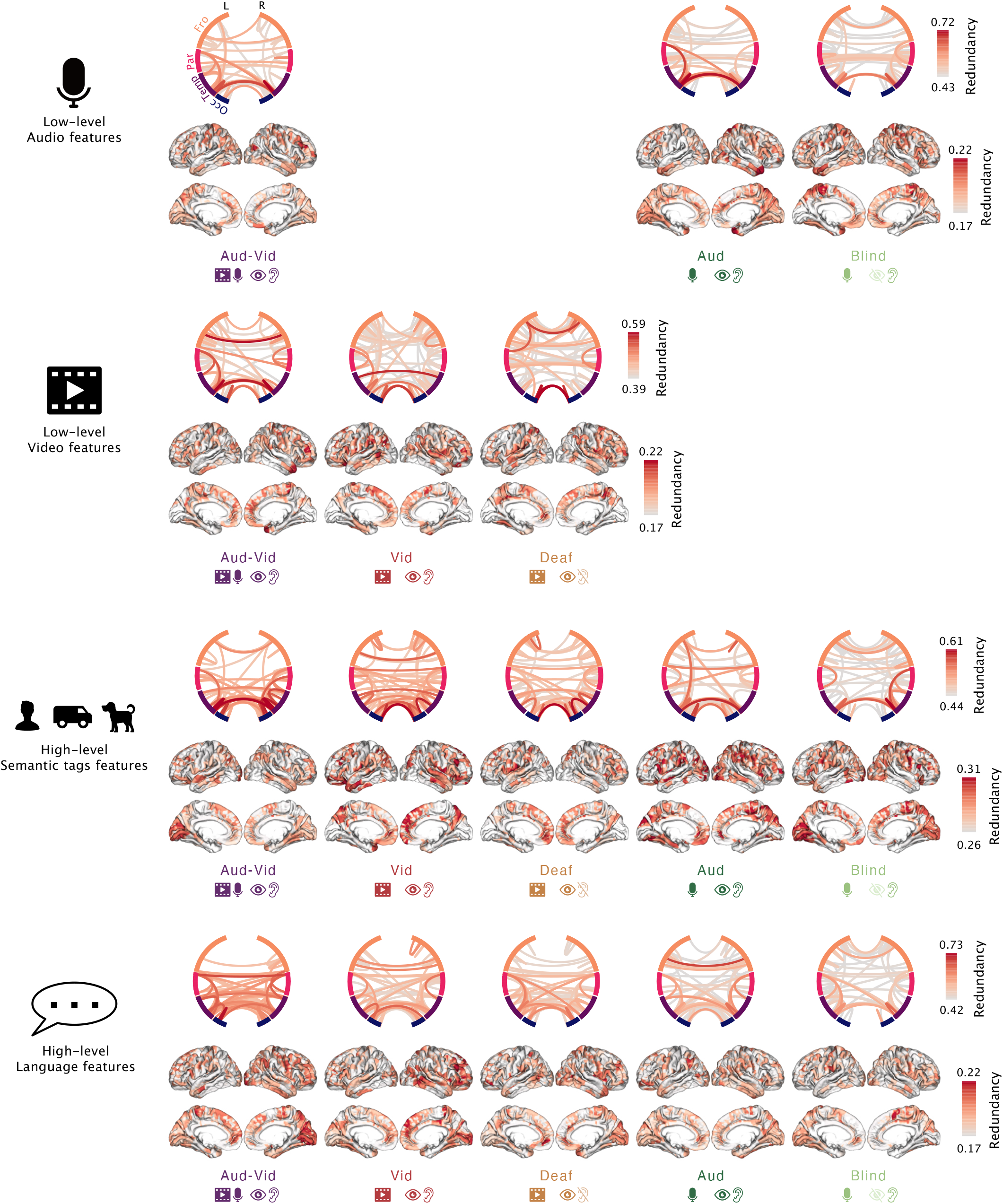
Redundant feature encoding across experimental groups and stimulus features. Group-average redundant information between pairs of cortical regions and stimulus features, estimated with Partial Information Decomposition (PID) for each experimental group and feature class. Circular plots show the top 100 redundant feature-encoding interactions, while brain maps show regional betweenness centrality of each cortical region based on the corresponding redundancy interaction profile. Rows correspond to low-level audio features, low-level video features, high-level semantic-tag features and high-level language features. These maps provide the full condition-wise distribution of redundant feature encoding underlying the PID analyses.

**Figure S6.**
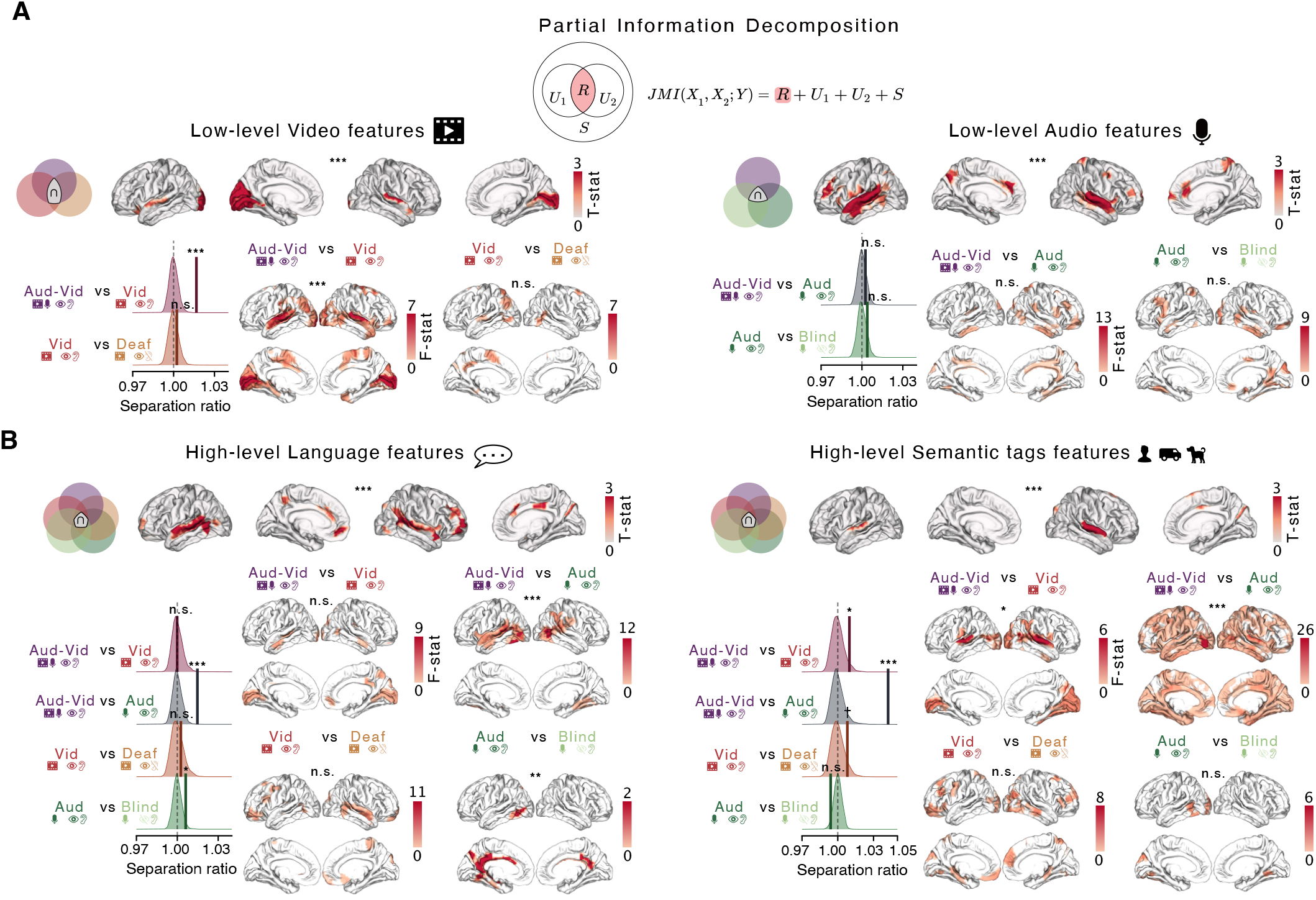
Redundant encoding of low-level and high-level perceptual stimulus features revealed by Partial Information Decomposition. **A-B**. For each stimulus feature encoding results, conjunction analyses (each top row) identify encoding patterns common to the relevant experimental groups, revealing cortical regions whose redundant encoding was invariant to sensory modality or sensory experience. Statistical significance was determined by cluster correction and permutation testing, while Venn diagrams indicate the groups included in each conjunction analysis. Separation-ratio plots show observed contrast statistics against permutation null distributions among contrasted whole-brain redundant interaction profiles. Brain maps show regions whose seed-to-whole-brain feature-encoding redundant interaction profiles differed between conditions, estimated using fcMVPA. Asterisks and transparency indicate statistical significance (P<0.05, cluster-corrected for brain plots). When fcMPVA showed no significant clusters, we highlighted regions with an F-statistic greater than 3. Legend: * P<0.05, ** P<0.01, *** P<0.001; n.s., not significant.

**Figure S7.**
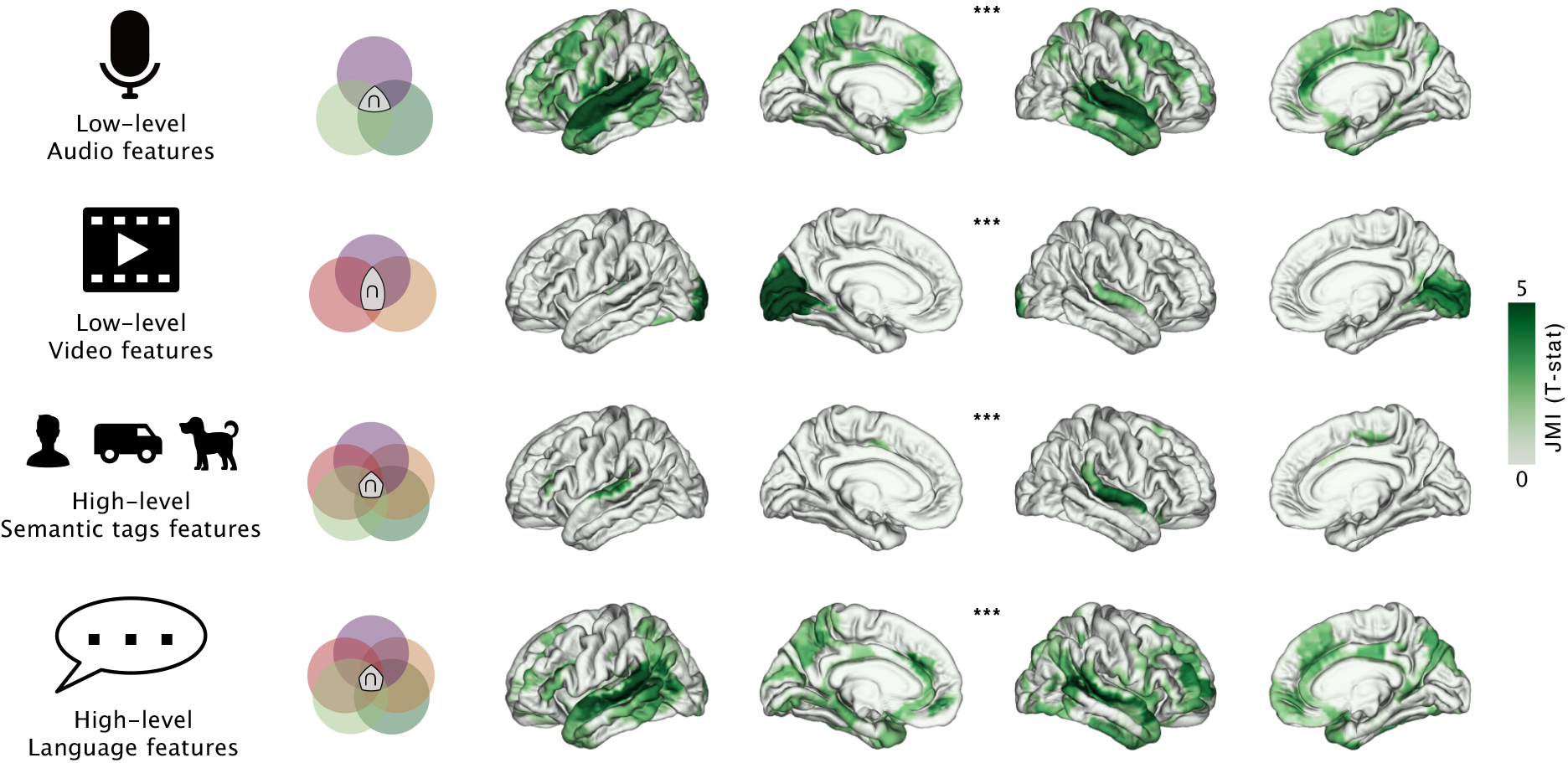
Conjunction analysis of joint mutual information across stimulus features. Conjunction analyses identifying cortical regions showing joint mutual information (JMI) common across the relevant experimental groups for each stimulus feature class. Rows correspond to low-level audio features, low-level video features, high-level semantic-tag features and high-level language features. Venn diagrams indicate the groups included in each conjunction analysis. Brain maps show conjunction t-statistics, highlighting cortical regions whose joint feature encoding was invariant to sensory modality and/or previous sensory experience. Asterisks indicate statistical significance determined by cluster correction and permutation testing (*** P < 0.001).

## Notes

### Competing Interest Statement

The authors have declared no competing interest.

